# A Proximity MAP of RAB GTPases

**DOI:** 10.1101/2024.11.05.621850

**Authors:** Véronique Gaudeault St-Laurent, Benoit Marchand, Raphaëlle Larcher, Sonya Nassari, Francis Bourassa, Mathilde Moreau, Dominique Jean, François-Michel Boisvert, Marie A. Brunet, Steve Jean

## Abstract

RAB GTPases are the most abundant family of small GTPases and regulate multiple aspects of membrane trafficking events, from cargo sorting to vesicle budding, transport, docking, and fusion. To regulate these processes, RABs are tightly regulated by guanine exchange factors (GEFs) and GTPase-activating proteins (GAPs). Activated RABs recruit effector proteins that regulate trafficking. Identifying RAB-associated proteins has proven to be difficult because their association with interacting proteins is often transient. Recent advances in proximity labeling approaches that allow for the covalent labeling of neighbors of proteins of interest now permit the cataloging of proteins in the vicinity of RAB GTPases. Here, we report APEX2 proximity labeling of 23 human RABs and their neighboring proteomes. We have used bioinformatic analyses to map specific proximal proteins for an extensive array of RAB GTPases, and RAB localization can be inferred from their adjacent proteins. Focusing on specific examples, we identified a physical interaction between RAB25 and DENND6A, which affects cell migration. We also show functional relationships between RAB14 and the EARP complex, or between RAB14 and SHIP164 and its close ortholog UHRF1BP1. Our dataset provides an extensive resource to the community and helps define novel functional connections between RAB GTPases and their neighboring proteins.

## INTRODUCTION

Cells actively communicate with their neighbors and adapt to signals and stress^1^. Membrane trafficking events are intrinsically linked to these responses^2,3^ by controlling their location and duration^4,5^. Therefore, they must be tightly regulated^6^. RAB GTPases are an essential class of proteins that regulate membrane trafficking events^7,8^.

RAB GTPases represent the largest family of small GTPase, with approximately 70 members in humans^9,10^. They are thought to act as molecular switches and cycle between active GTP-bound and inactive GDP-bound states^11^. Rapid cycling is facilitated by guanine exchange factors (GEFs) and GTPase-activating proteins (GAPs)^12,13^. Once activated, RABs interact with a wide range of effector proteins^14^ at specific membrane locations^15^ to regulate several aspects of vesicular trafficking. They can interact with coat proteins to regulate cargo sorting^16^, with molecular motors to control vesicle movement^17,18^, and with tether^19,20^ and SNAREs to modulate vesicle fusion^21^. RAB GTPases can also directly control signaling events, with RAB35 acting as an oncogene when mutated by constitutively activating the PI3K/AKT pathway^22^.

RABs can undergo RAB cascades to ensure the appropriate progression of endocytosed or secretory cargos^8^. A well-described example is the RAB5 to RAB7 transition regulated by the binding of RAB7 GEF Mon1/CCZ1 by RAB5^23,24^ and the recruitment of RAB5 GAP TBC1D18 by Mon1^25^. Interestingly, these RAB cascades often coincide with phosphoinositide transitions^15^. Finally, although RABs are primarily associated with specific membrane compartments, they can be recruited to ‘non-conventional’ sites under specific stimuli or stresses. An example of this is RAB5 translocation to the mitochondria upon oxidative stress^26^.

In addition to regulation by GEFs and GAPs, post-translational modifications of RABs can directly affect their capacity to bind to effectors or cycle^27,28^. RABs phosphorylation^29,30^ and palmitoylation^31^ also directly modulate their function. Pathogens have evolved to co-opt or inhibit RAB GTPases through post-translational modifications^32^. These examples highlight the numerous roles that RABs play, their intricate regulation, and their capacity to interact with regulators and effectors depending on the cellular context. Defining the scope and dynamics of the interacting proteins is essential for understanding the function of RAB GTPases.

Much progress has been made recently in defining RAB GTPases direct and proximal interactors. Earlier studies have used yeast two-hybrid^33^ or *in vitro* affinity purifications^34,35^ to identify direct RAB-binding proteins. These methodologies performed well in identifying potent binding effectors and regulators, but did not allow for the interrogation of RAB interactors under specific cellular conditions. Lee *et al*^36^ performed direct affinity purification of 51 TAP-tagged RABs to build a RAB interactome. This effort led to the identification of an extensive array of potential RAB-interacting proteins, with the caveat that dual TAP-tag purification led to the absence of most of the known interactors for various RABs^36^. The advent of proximity ligation approaches^37^ and BioID has dramatically changed the ability to determine protein neighbors of tagged proteins^38^. Initially, BioID used a mutated *E. coli* biotin ligase (BirA) that allowed for efficient proximity labeling over a 24 h period^37^. Directed evolution yielded enhanced BioID enzymes^39–41^, providing a shorter labeling time and allowing for the investigation of dynamic protein interactors or localization^42^. Similarly, the peroxidase APEX2 has been developed for electron microscopy and proximity labeling^43^ and gives, through the addition of hydrogen peroxide, a snapshot of proximal proteins of a bait of interest^44,45^. All BioID approaches have the caveat that the enzymes used are ≥ 19 kDa in size and can potentially interfere with protein function. Furthermore, the labeling radius varies among enzymes, with APEX2 being associated with a greater background, especially in cytosolic proteins^46^. Nevertheless, BioID methodologies have allowed for many discoveries and a better understanding of protein localization and dynamics^47,48^. BioID was used to study small GTPases. As such, we previously relied on APEX2-mediated BioID of three early endosomal RABs to demonstrate a novel link between RAB21 and endosomal sorting complexes^49^, while Gillingham et al. targeted GDP or GTP-locked small BirA-tagged RAB GTPases to the mitochondria^50^ and proved the usefulness of this approach for mapping regulators and effectors. Dynamic APEX2 studies were also performed on RAB5^51^ and TBC1D15^52^ to determine stimuli-specific interactions. Large-scale studies on Rho GTPases^53^ and Arf GTPases^54,55^ further illustrate the power and usefulness of comparing multiple prey species to identify specific and novel regulators from background interactions.

In this study, we generated a broad proximal RAB interactome using APEX2-tagged RABs. We report that APEX2:RAB fusion protein localization can be predicted from identified neighboring proteins. We demonstrate novel interactions between multiple RABs and establish a relationship between RAB25 and DENND6A during cell migration. Furthermore, we show that RAB14 is required for endosomal recruitment of the EARP complex. We used our dataset to show a functional link between RAB14 and the lipid transfer proteins SHIP164 and UHRF1BP1.

## RESULTS

### Most APEX2:RABs localize to their expected site and biotinylate protein neighbors

We had previously identified a new interaction between RAB21 and the WASH complex through APEX2-mediated proximity ligation of three early endosomal RAB GTPases: RAB4a, RAB5a, and RAB21^49^. Given the utility of APEX2:RAB fusions to interrogate proteins in the vicinity of RAB GTPases^49,56^ (Fig. 1A), we aimed to generate a larger APEX2:RAB proximity map. Therefore, we generated fusion proteins for 24 human RABs localized to different membrane compartments (Fig. 1B) and distributed throughout the RAB evolutionary tree^29^ (Suppl. Fig. 1A). Flp-in T-Rex stable cell lines were generated and most RABs were expressed as full-length proteins (Suppl. Fig. 1B). These proteins localized to their expected compartments, and led to local biotinylation in those compartments (see RAB1A and RAB2A Golgi localization), while RAB35 was enriched at the plasma membrane (Fig. 1C). Notably, despite appropriate cloning, a few APEX2:RABs (RAB8, 13, 20, 28, 29, and 34) were not expressed at detectable levels (Suppl. Fig. 1B). Finally, most APEX2:RABs that were expressed, were able to biotinylate neighboring proteins upon biotin-phenol addition and H_2_O_2_ treatment, except for RAB24 and RAB31. (Suppl. Fig. 1B). The lack of biotinylation for these two RABs could be caused by their enriched localization at mitochondria or peroxisomes^47^ or lower expression levels (Suppl. Fig. 1B). From these results, we conclude that the APEX2:RAB stable cell line library is a good resource for performing mass spectrometry (MS) analysis of RAB proximal proteins.

**Figure 1:**
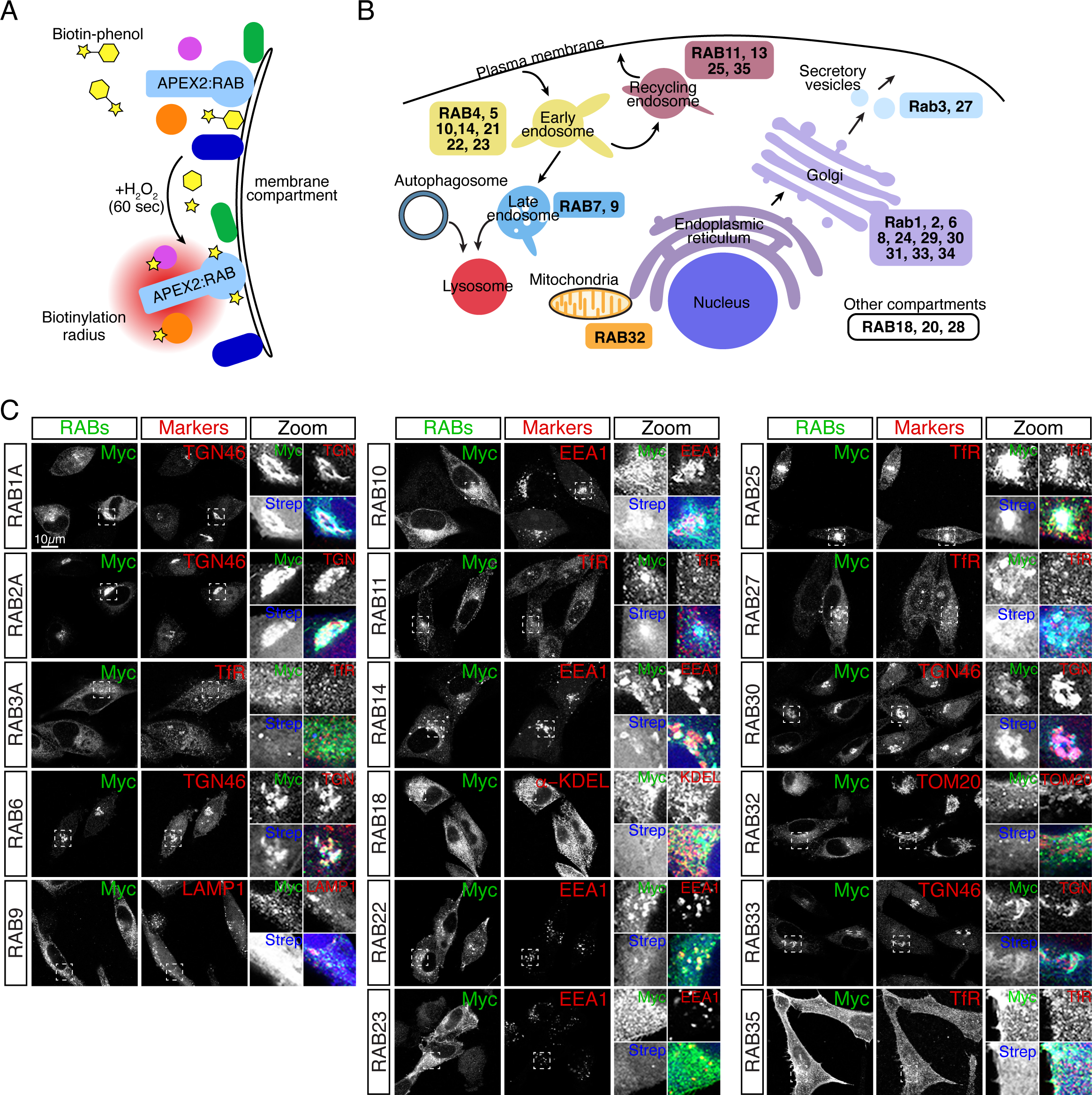
APEX2:RABs appropriately localize and lead to local biotinylation. Cartoon representation of **(A)** APEX2-mediated proximity labeling and **(B)** primary localization of the RABs assessed in this study. **(C)** Immunofluorescence analysis of APEX2:RABs (MYC), specific organelle markers (TGN46, Transferrin receptor (TfR), EEA1, LAMP1, and TOM20) and biotinylated proteins (streptavidin). Boxes highlight zoomed views of all stainings and merged images, *n*=3 independent experiments. Scale bars: 10µm. Note that RAB4A, RAB5A, RAB7A and RAB21 localizations and abilities to biotinylate neighboring proteins were published in a previous study by our group^49^. Hence, we did not add validation experiments in this figure.

### APEX2:RAB neighboring proteins are associated with membrane trafficking processes

(MS) analyses were performed in triplicates for each RAB protein, and statistical enrichment analysis of the dataset was performed using SAINT^57^. APEX2:RABs self-biotinylation was detected at a high level and was statistically significant for most RABs except for RAB9A (Suppl. Fig. 1C). The different degrees of self-biotinylation could be caused by the types of residues exposed in the various RABs, as activated biotin reacts more potently with tyrosines^58^. Analysis of high-confidence hits (Extended Table 1) identified for each RAB highlighted significant differences in the number of observed preys between RABs (Fig. 2A). For example, early endosomal RABs averaged 1087 significantly enriched proximal proteins, with an average of 35 for late endosomal RABs. These differences could be due to the availability of the biotin-phenol in various compartments populated by diverse RABs. Nonetheless, by comparing individual RAB to the human CellMap dataset^47^ generated with BirA-mediated proximity labeling, we could successfully infer APEX2:RAB localization for most RABs (Suppl. Fig. 2A). This analysis highlighted some discrepancies between the expected and inferred localizations of four RABs. The RAB11, 22, 23, and 27 datasets emphasized nuclear localization (Suppl. Fig. 2A), which is unlikely, and probably driven by their proximity to splicing factors (Extended Table 1). We performed GO term enrichment analyses using Metascape^59^. For most RABs, membrane trafficking processes were identified with high confidence (-log10(P) values lower than 10^-15^ for these terms). Again, the RAB11, 22, 23, and 27 datasets showed strong enrichment for nucleus-related processes (Suppl. Fig. 2B). Therefore, we excluded these RABs from further analyses. By analyzing the proximal proteins for the remaining RABs we observed the enrichment of highly significant GO terms linked to vesicular processes (Fig. 2B). Network-based representation of these GO terms highlighted the tight connectivity between the terms, further strengthening the usefulness of the generated dataset (Fig. 2C).

**Figure 2:**
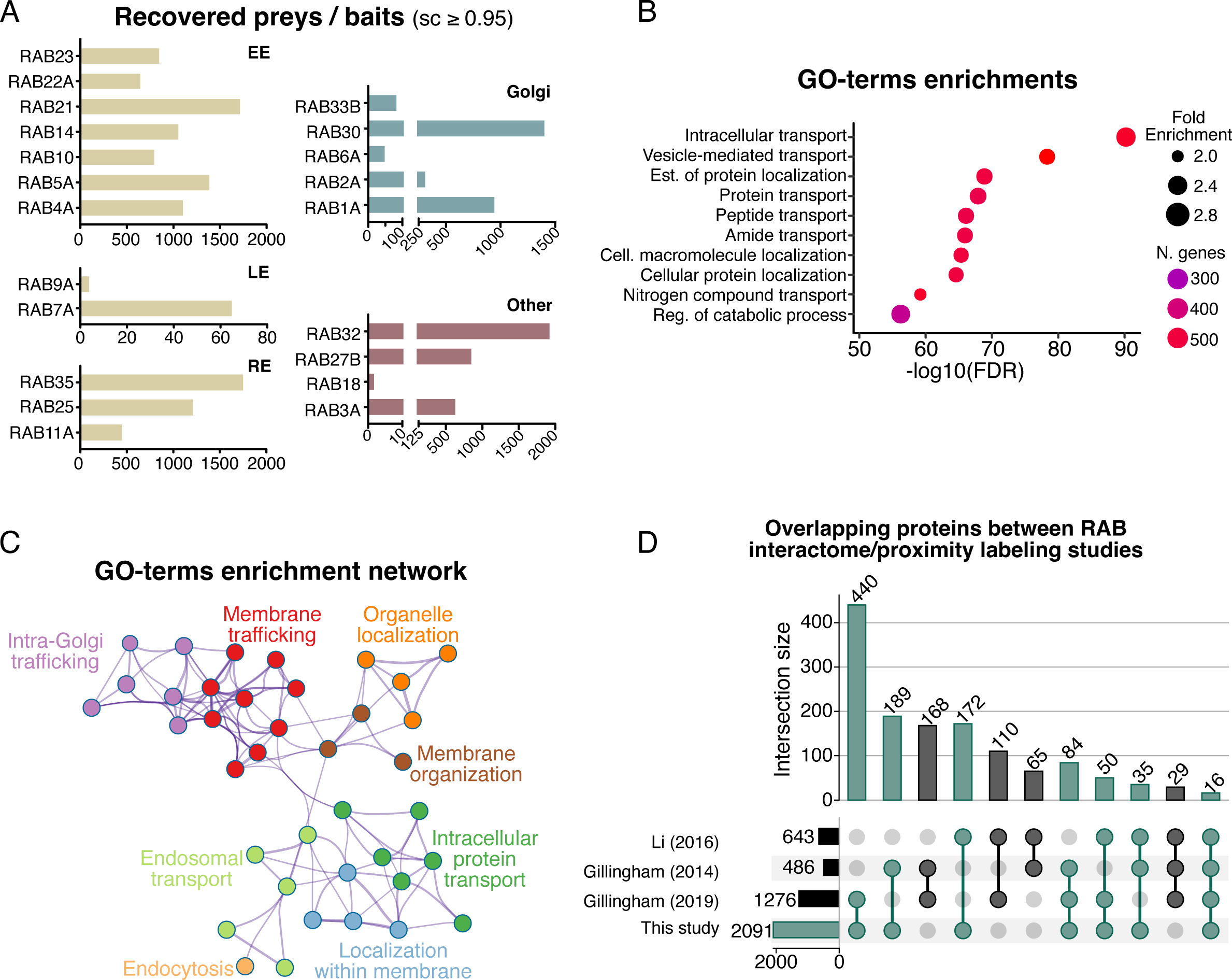
Global analysis of the APEX2:RAB-generated proximity map. **(A)** Number of identified preys per RAB. RABs were separated based on their general localizations. Previously published RAB4A, RAB5A, RAB7A and RAB21 data was incorporated in the general proximity interactome map. **(B)** Go-terms enrichment analysis of all proteins identified with a SC above 0.95. Note that RAB11, 22, 23, and 27 datasets were excluded. **(C)** Network representation of enriched ontology clusters linked to membrane trafficking events. Colors represent clusters and each node represents a GO-term. **(D)** Upset plots of three RAB interactome studies compared to the new APEX2:RAB dataset presented in this study, detailing the number of overlapping proteins.

Additionally, we compared our APEX2 proximity map with those of other RAB GTPase interactomes^35,36,50^. As not all RABs were tested in the four studies, we compared the data for RAB2, 4, 5, 6, 7, 9, 10, 11, 18, and 32. We combined data from various studies in which GTPases were analyzed in their GDP- or GTP-bound states including all paralogs if they were tested (i.e., RAB5a, RAB5b, and RAB5c). Approximately 35% of the identified proteins in each interactome were recovered from our dataset (35% Gillingham et al. 2019; 39% Gillingham et al. 2014 and 26% Li et al. 2016) (Fig. 2D). When the number of overlaps was compared between multiple studies, the overlap between the identified proteins decreased (Fig. 2D). It is worth noting that the highest number of overlapping proteins was found between the mitoID-generated proximity interactome (Gillingham, 2019) and our dataset, probably because of the higher number of identified proteins and similarity of the experimental design. Interestingly, the highest similarity in GO-terms enrichment was observed with proximity labeling protocols (Suppl. Fig. 2C). At the same time, the overlap was minimal when the interactomes generated from immunoprecipitations (Li, 2016) and GST-affinity purification (Gillingham, 2014) were compared (Suppl. Fig. 2C). Combining these analyses, we conclude that the APEX2:RAB proximity interactome is specifically enriched, to varying degrees, for proteins associated to RAB-linked processes and offers a resource for identifying new functional relationships between RAB GTPases and their neighboring proteins.

### Known RAB regulators can be observed, and novel relationships can be extracted from the APEX2:RAB dataset

BioID approaches enable the identification of large arrays of proteins, making it challenging to identify specific interactors^38^. We used two scores to identify novel relationships between RABs and their regulators. We utilized a specificity score (WD score)^60^ developed to identify neighboring proteins that were enriched more specifically in a given bait (RABs in this case). Using this score and selecting statistically significant known and predicted GEFs and GAPs, we identified known RAB-GEF and RAB-GAP pairs (Suppl. Fig. 2D and E). As examples, and as detected previously, VARP, a RAB21 GEF, showed the highest specificity for RAB21^49^, while it was also observed with RAB32 (Suppl. Fig. 2D), a known VARP interactor^61^. The RGP1-RIC1 heterodimer acts as an RAB6 GEF^62,63^. Our data revealed that RAB6A showed a strong specificity for RGP1 (Suppl. Fig. 2D), whereas RIC1 was observed in RAB6A dataset, but to a threshold inferior to our exclusion criteria (Fig. 3A). RIC1 was proximal to several other RABs, particularly RAB30. Hence, it is possible that RIC1, in addition to its GEF activity towards RAB6A, has some functional links with RAB30. Known RAB GAP pairs were also highlighted in the proteomic dataset. TBC1D10B and RN-TRE (USP6-NL), which have RAB35 GAP activity^64^, were specific for RAB35 (Fig. 3B and Suppl. Fig. 2E). Interestingly, the RAB11 GAP TBC1D8B was highly specific for RAB25, an ortholog of RAB11^65^ (Suppl. Fig. 2E). Additionally, a BioID study of recycling endosomes RABs^66^ identified TBC1D2B as a RAB25 proximal protein (Suppl. Fig. 2E). Some RAB GEFs and GAPs have been found to be associated with multiple RABs. GAPEX5 was observed to be associated with most RABs, which could be explained by its activity on RAB5 and RAB31 and its association with clathrin-coated and microtubule regulatory proteins^67,68^, leading to its enrichment in the vicinity of various RABs. TBC1D5, which is a well-characterized RAB7A GAP, displayed the highest specificity with RAB7A^69^ but was also observed in proximity of other RABs (Suppl. Fig. 2E). TBC1D5 localization is associated with numerous cellular compartments^70,71^, potentially leading to its enrichment in various baits.

**Figure 3:**
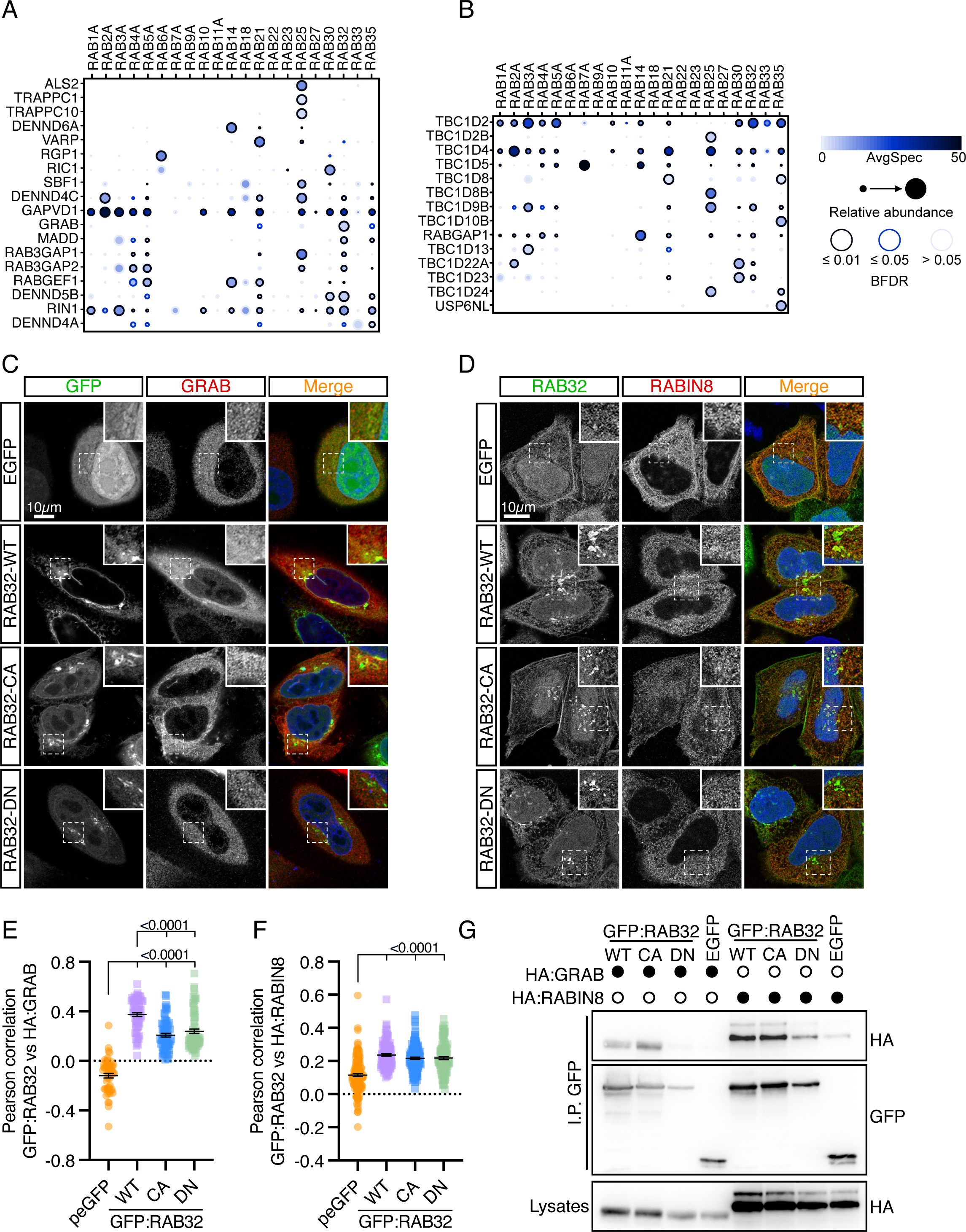
Analysis of RABs proximity to GEFs and GAPs identifies novel connections. **(A and B)** Prohits-viz generated dotplots for (A) GEFs and (B) GAPs identified enriched with a Saint score ≥ 0.95 for all the RABs tested in this study. **(C and D)** RAB32 partially colocalizes with (C) GRAB and (D) RABIN8. Transiently expressing GFP, GFP:RAB32-WT, Q85L (CA) and T39N (DN) and HA:GRAB or HA:RABIN8 HeLa cells were fixed and stained. Boxed regions are magnified, and single channels are depicted. Scale bars: 10µm. **(E)** Pearson’s correlation between RAB32 and GRAB. Error bars are SEM, individual points represent single cells, *n*=3 independent experiments. **(F)** Pearson’s correlation between RAB32 and RABIN8. Error bars are SEM, individual points represent single cells, *n*=3 independent experiments. **(G)** RAB32 interacts with GRAB and RABIN8. GFP-Trap IP of RAB32 WT, CA, and DN variants in HeLa cells followed by GFP and HA immunoblots, *n=3* independent experiments. Stats performed in (E) and (F) were Kruskal-Wallis followed by a Dunn’s post test.

We also relied on dot plot representations^57^ that displayed relative abundance and statistical significance compared to the control condition in a simple graphical representation (Fig. 3 A and B). Using this graphical representation, strong enrichment of ALS2 and TRAPPC1/C10 with RAB25 was evident (Fig. 3A). Interestingly, abundance of VARP was higher in association with RAB21, but was also significantly detected with RAB32 (Fig. 3A). Similar results were observed for DENND6A, which showed the highest specificity and abundance with RAB14, its known GEF^72^ (Fig. 3A and Suppl. Fig 2D). Finally, GRAB, which is a close paralog of RABIN8 and acts as a RAB8 GEF and RAB11 effector^73^, showed specificity towards a few RABs (Suppl. Fig. 2D). However, its relative abundance was higher with RAB32 (Fig. 3A). We focused on RAB32 and GRAB as well as RAB25, ALS2, and DENND6A proximities to confirm potential new relationships. These pairs were selected based on their specificity scores and statistically significant enrichments (Fig. 3A and S2D). Since GRAB is a close RABIN8 paralog with known activity towards RAB8, we performed co-localization and interaction analyses between WT, GTP-bound constitutively active (CA), and GDP-bound dominant negative (DN) RAB32. Colocalization studies revealed that GRAB and RABIN8 displayed a cytosolic localization pattern with limited colocalization with RAB32 (Fig. 3C–F), except for wild-type RAB32, which displayed higher colocalization (Fig. 3C and E). Co-immunoprecipitation experiments between GRAB and RABIN8 demonstrated an interaction of both with wild-type and activated RAB32 (Fig. 3G). Proteomic proximity and validation of the interaction among RAB32, GRAB, and RABIN8 indicates that RAB32 could act as a novel GRAB or RABIN8 effector.

### RAB25 interacts with ALS2 and DENND6A, and its localization is affected by DENND6A

RAB25 is an atypical RAB with poor GTPase activity^74^ and is involved in various oncogenic processes^75–78^. Given its proximity to various GEFs, we aimed to validate these observations and test their functional relationships. The TRAPPC complex acts as RAB1 and RAB11 GEFs but has no activity towards RAB25^79–81^. Since RAB25 is a member of the RAB11 family, the proximity we observed could be related to a potential effector function of RAB25 with TRAPP. Additionally, we observed a high relative abundance between RAB25 and ALS2 a RAB5 GEF (Fig. 3A). Furthermore, we observed the proximity with lower specificity and relative abundances of multiple GEFs with RAB25. DENND6A, a known RAB14 GEF^72^, was of interest because it is poorly characterized, showed some specificity, and was significantly enriched in RAB25 proximal proteins (Fig. 3A and Suppl. Fig. 2D). Therefore, we assessed the colocalization between RAB25, ALS2, and DENND6A. We used MCF7 cells since they express RAB25 (Suppl. Fig. 3D). GFP:RAB25 was localized to intracellular puncta, whereas ALS2 displayed a mostly cytosolic localization pattern (Fig. 4A). ALS2 did not extensively colocalize with RAB25, as evidenced by its low Pearson colocalization coefficient (Fig. 4B). Nevertheless, we confirmed the MS data and coimmunoprecipitated ALS2 with RAB25 (Fig. 4C). Collectively, these data highlight a potential functional link between ALS2 and RAB25.

**Figure 4:**
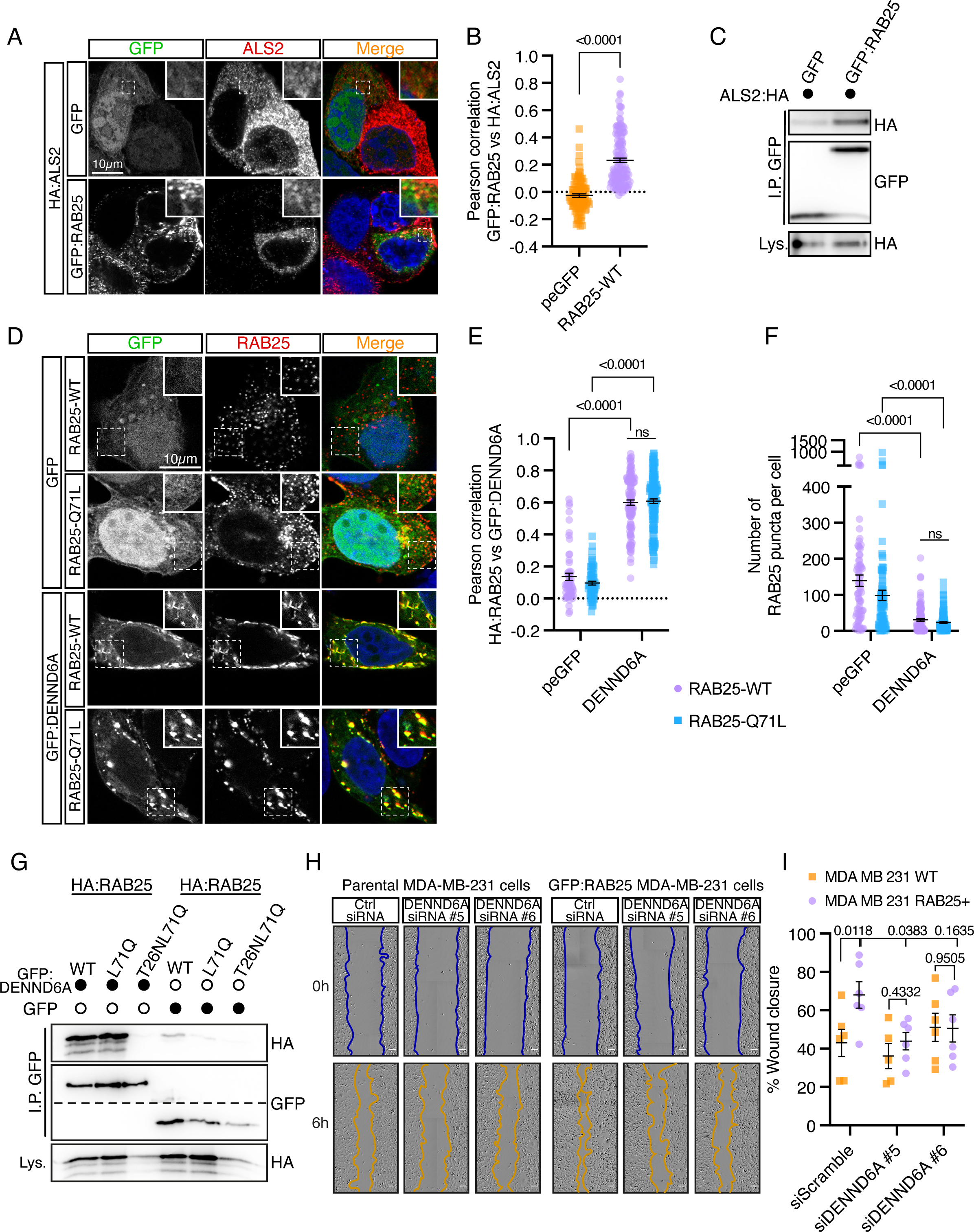
RAB25 interacts physically and functionally with DENND6A. **(A)** RAB25 partially colocalizes with ALS2. Transiently expressing GFP, GFP:RAB25 and HA:ALS2 MCF7 cells were fixed and stained. Boxed regions are magnified, and single channels are depicted. Scale bars: 10µm. **(B)** Pearson’s correlation between RAB25 and ALS2. Error bars are SEM, individual points represent single cells, *n*=3 independent experiments. **(C)** RAB25 interacts with ALS2. GFP-Trap IP of GFP or GFP:RAB25 in MCF7 cells followed by GFP and HA immunoblots, *n=3* independent experiments. **(D)** RAB25 relocalizes in DENND6A overexpressing cells. Transiently expressing HA:RAB25 WT or Q71L MCF7 cells coexpressing GFP or GFP:DENND6A were fixed and stained. Boxed regions are magnified. Scale bar: 10µm. **(E)** Pearson’s correlation between RAB25 and DENND6A. Error bars are SEM, individual points represent single cells, *n*=3 independent experiments. **(F)** Quantification of RAB25 puncta per cell. Error bars are SEM, individual points represent single cells, *n*=3 independent experiments. **(G)** RAB25 interacts with DENND6A. GFP-Trap IP of RAB25 WT, L71Q, and T26NL71Q variants or GFP in MCF7 cells followed by GFP and HA immunoblots, *n=3* independent experiments. **(H)** The RAB25-mediated increased migration of MDA-MB-231 cells is blunted by DENND6A knockdown. Brightfield images of wounded MDA-MB-231 monolayer at the beginning of the experiment (top) and 6 hours after wounding (bottom). **(I)** Percentage of wound closure 6 hours after wounding. Error bars are SEM, individual points represent single experiments, *n*=6 independent experiments. Stats performed were (B) Two-tail Mann Whitney test, (E and F) 2-way ANOVA with Sidak post test and (I) 2-way ANOVA with Tukey post test.

Although RAB25/DENND6A proximity was less abundant and specific than ALS2/RAB25 (Fig. 3A), wild-type RAB25 and DENND6A extensively colocalized on intracellular puncta in MCF7 cells (Fig. 4D and E). Given this high colocalization, we tested if restoring the cycling ability of RAB25 or locking it with GDP would affect its colocalization with DENND6A. We expressed WT (which for RAB25 is mainly associated with GTP), RAB25Q71L (which is equivalent to a WT GTPase and can cycle), and RAB25-T26NQ71L (a dominant negative GTPase, GDP-bound) with DENND6A, and tested for colocalization. We found that cycling (Q71L) did not reduce colocalization (Fig. 4D and E). However, the expression of T26NQ71L RAB25 resulted in diffuse RAB25 localization (Suppl. Fig. 3A). Hinting at a potential functional relationship between DENND6A and RAB25, we observed that DENND6A expression drastically affected RAB25 localization. While reduced in number, RAB25-positive individual puncta were enlarged and highly colocalized with DENND6A-positive puncta (Fig. 4D and F). Furthermore, no differences in colocalization were observed between DENND6a and RAB14, a well-described target, in RAB25 knockout MCF7 cells compared to WT (Suppl. Fig. 3B, C and D). This result suggests that DENND6A is more likely to modulate RAB25 localization than the opposite. Furthermore, earlier this year, a study confirmed our findings by demonstrating that mislocalized mito-DENND6A increased the translocation of RAB25 to the mitochondria^82^. In accordance with the colocalization studies, co-immunoprecipitation experiments showed that DENND6A interacted with both the WT and RAB25 Q71L variants but not with the GDP-locked form (Fig. 4G). Together, these results demonstrate the usefulness of proximity labeling proteomic data in identifying novel interactions between RAB GTPases and GEFs.

### DENND6A affects RAB25-driven cell migration

Given the modulation of RAB25 localization by DENND6A, we investigated whether DENND6A modulates RAB25-driven processes. Overexpression of RAB25 improves cell migration on the extracellular matrix^75^. Therefore, we generated stable MDA-MB-231 cells overexpressing GFP:RAB25 (Suppl. Fig. 3E) and wound healing assays were performed using fibronectin-coated dishes. As observed in ovarian cancer cells^75^, RAB25 expression accelerated wound closure (Fig. 4H and I). Moreover, RAB25 expression in MDA-MB-231 significantly increased the expression of *DENND6A* (Suppl. Fig. 3F). Interestingly, the depletion of DENND6A expression by siRNA (Suppl. Fig. 3G) did not affect wound closure in the parental MDA-MB-231 cells (Fig. 4H and I). However, it only slightly, but significantly, reduced wound closure in MDA-RAB25 cells (Fig. 4H and I). These results suggest a relationship between DENND6A and RAB25, where RAB25 expression increases the expression of DENND6A required for its adequate localization to sustain an increased cell migration. These turns of events could be related to the RAB25-dependent increased expression of CLIC3, a protein required for the recycling of α5β1 integrins routed to the lysosome by RAB25, a recycling pathway known to increase cancer cell migration efficiency^77^. RAB25 plays well-defined roles in integrin recycling^77,83^. Given the migratory phenotype, we analyzed the levels and localization of total (Suppl. Fig. 3G and H), and active ≥1 integrin (Suppl. Fig. 3I and J) in parental or MDA-RAB25 cells with or without DENND6A depletion. No significant consistent differences were observed (Suppl Fig. 3H-K), indicating that the observed effects were unrelated to integrin expression or activation levels. Altogether, these experiments established novel interactions between RAB32 and GRAB/RABIN8, as well as with RAB25/ALS2 and RAB25/DENND6A.

### APEX2 proteomic identifies proximities between RAB GTPases and trafficking complexes

RABs modulate specific trafficking events by interacting with membrane-associated complexes^7^. A prime example is the capacity of RAB7A to recruit the retromer complex to modulate endosomal cargo sorting events^84^ or the interaction between RAB11 and the exocyst complex to modulate exocytosis^85^. Additionally, RAB5 and RAB7 can interact with Corvet and HOPS complexes to modulate early and late endosome fusion events, respectively^86,87^. To identify novel relationships between RABs and sorting complexes, we analyzed the proximity between RABs and membrane trafficking regulatory complexes. As shown previously^49^, we observed proximity between RAB7A and the retromer with the highest specificity (Fig. 5A and suppl. Fig. 4A). Multiple subunits of the Corvet complex were specific to RAB5. In contrast, RAB25 showed proximity to most exocyst subunits, with the highest specificity (Suppl. Fig. 4A). This finding is consistent with the role of RAB25 in apical recycling^88^. Interestingly, we observed a strong specificity and abundance between COG complex members and RAB2A (Fig. 5A and Suppl. Fig. 4A), which is consistent with the results of yeast two-hybrid interaction analysis between RAB2 and COG5^89^. Our MS data suggested a higher proximity between the COG complex and RAB2, followed by RAB30, RAB1, and RAB25, showing some proximity but with a lower confidence level (Fig. 5A). We tested the proximity between RAB2 and COG1, COG4, and COG6 using proximity ligation assays and observed a high proximity to RAB2 (Suppl. Fig. 4B and C).

**Figure 5:**
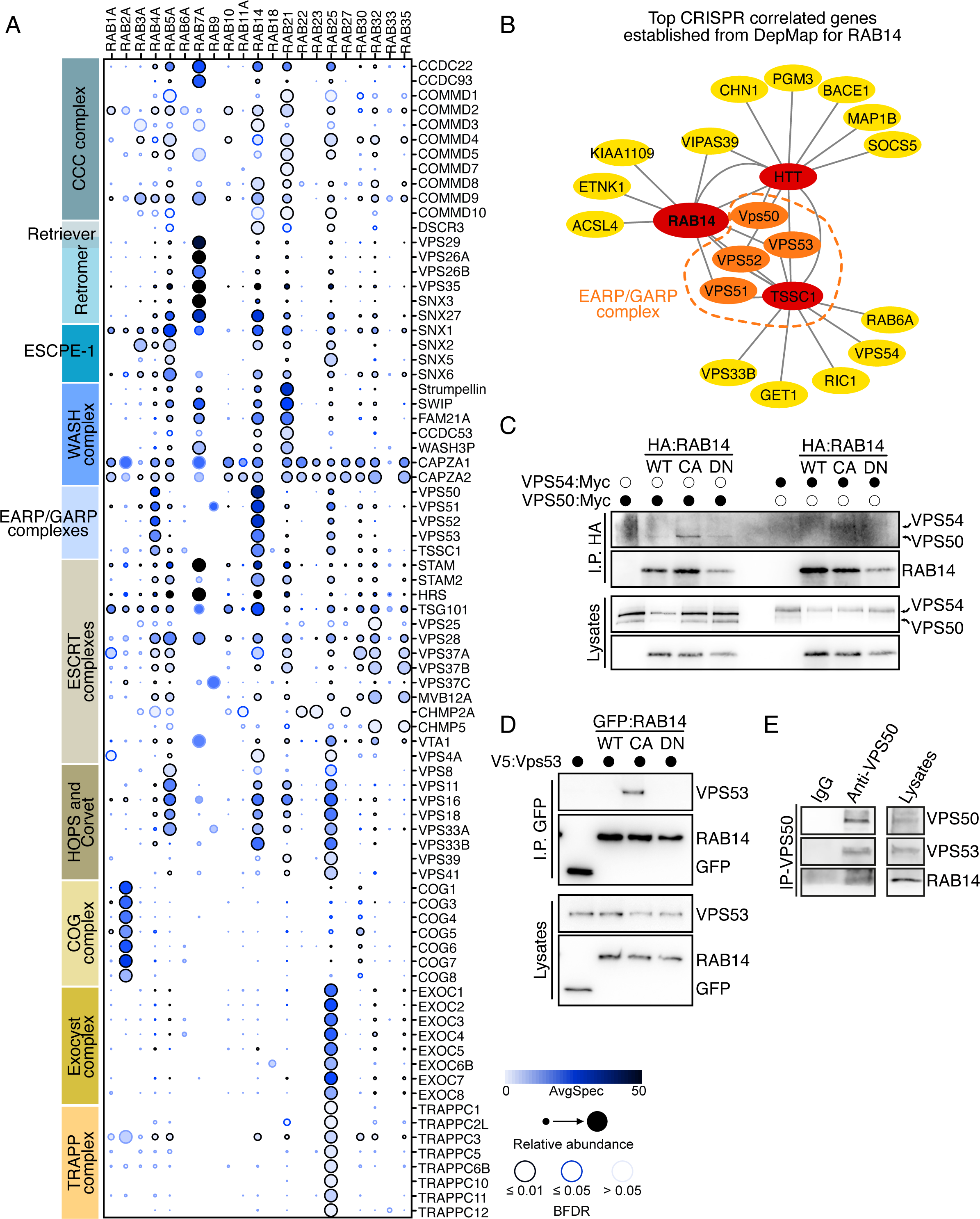
RAB14 interacts with members of the EARP complex. **(A)** Prohits-viz generated dot plot for membrane trafficking complexes found enriched with a Saint score of ≥ 0.95 for all RABs tested in this study. **(B)** Top 10 most correlated genes with RAB14 in co-dependency screens from DepMap^92^. Red-colored nodes represent the top two correlated genes with RAB14, orange are EARP complex members, and yellow represents correlated genes with the three red nodes. **(C)** Activated RAB14 interacts with VPS50 and weakly with VPS54. GFP-Trap IP of RAB14 WT, Q70L (CA), and S25N (DN) variants or GFP in HeLa cells followed by GFP and MYC immunoblots, *n=3* independent experiments. **(D)** Activated RAB14 interacts with VPS53. GFP-Trap IP of RAB14 WT, Q70L (CA), and S25N (DN) variants or GFP in HeLa cells followed by GFP and V5 immunoblots, *n=3* independent experiments. **(E)** Endogenous VPS50 interacts with RAB14. Control IgG or anti-VPS50 rabbit antibody immunoprecipitations of HeLa cells lysates, followed by RAB14, VPS50 and VPS53 immunoblots. The cell lysate used for the control or VPS50 IPs was the same, *n=3* independent experiments.

The endosomal EARP complex^90^ stood out as proximal to RAB4 and RAB14, and its relative abundance was the highest with RAB14 (Fig. 4A). The interaction between RAB4 and EARP complex members has previously been observed^35^, but their interaction and functional links with RAB14 have not been tested. The EARP complex shares most of its subunits (VPS51, 52, and 53) with the Golgi-localized GARP complex^90,91^. Vps50 (CCDC132/Syndetin) is exclusive to the EARP complex, whereas VPS54 is specific to the GARP complex^90^. Interestingly, the VPS54 subunit was not observed in RAB4 or RAB14 proximity interactomes (Fig. 5A). To further justify the potential functional relationship between RAB14 and the EARP complex, we explored genetic correlations using the DepMap consortium^92^. Significantly, all subunits of the EARP complex, among the ten most correlated genes with RAB14, were identified (Fig. 5B, orange). We performed proximity ligation and co-immunoprecipitation studies between VPS50:MYC or VPS54:MYC and wild type, constitutively active (CA), and dominant-negative (DN) RAB14 to validate the MS data. Proximity labeling revealed a statistically significant proximity between VPS50 and all RAB14 forms, while VPS54 was also proximal to WT and CA RAB14 (Suppl. Fig. 5A and B). Proximity between RAB14 WT and VPS50 or VPS54 was significantly higher than that observed for dominant-negative RAB14 (Suppl. Fig 5A and B). Furthermore, supporting the lack of VPS54 in the MS data, RAB14-WT showed significantly more PLA puncta with VPS50 than with VPS54 (Suppl. Fig. 5A and B). Together, these results demonstrate the proximity between WT or active RAB14, with a preference for VPS50 over VPS54, and thus, showing more proximity with the EARP complex over the GARP complex. We validated the PLA results using co-immunoprecipitation analysis and observed an interaction between VPS50 and activated RAB14 (Fig. 5C). We also confirmed an interaction between RAB14 and VPS53, another subunit of the EARP complex (Fig. 5D). Significantly, we could recover endogenous RAB14 following the immunoprecipitation of endogenous VPS50 from HeLa cell lysates (Fig. 5E). This demonstrates that the Rab14 interaction with the EARP complex is observed endogenously in cells. We conclude from these results that RAB14 interacts with the EARP complex, which is consistent with findings from *C. elegans* that identified a genetic relationship between the EARP complex and Rab14^93^ and with previously published proteomic data^35^. This interaction also favors the activated form of RAB14, suggesting that the EARP complex could act as an RAB14 effector.

### RAB14 regulates EARP endosomal recruitment and impacts SNARE localization

Previous studies have independently implicated RAB14 and the EARP complex in transferrin receptor recycling^72,90^. Given the proteomic and co-immunoprecipitation results, we tested whether RAB14 could be involved in EARP complex endosomal recruitment. We generated clonal RAB14 knockout HeLa cell lines. The RAB14 knockout was validated by sequencing and western blot analysis (Suppl. Fig. 5C). We relied on VPS50 or VPS54 overexpression because commercial antibodies did not allow for specific labeling. Using MYC-tagged VPS50 and VPS54^90^, we detected co-localization between VPS50 or VPS54 and the transferrin receptor on intracellular puncta or in Golgi-like structures, respectively (Fig. 6A-C). Significantly, the RAB14 knockout led to the dispersion of VPS50 and reduced its colocalization with the transferrin receptor (Fig. 6A and B). No differences in VPS54 localization pattern or colocalization with the transferrin receptor were observed in RAB14-deleted cells (Fig. 6A and C). Overall, these observations support the requirement of RAB14 for endosomal recruitment of the EARP complex in HeLa cells. Since the EARP complex acts as an endosomal tethering complex^90^, we tested whether the loss of RAB14 affected the localization of EARP-associated SNARE^90^, by comparing the impact of RAB14, EARP (VPS50) (Suppl. Fig. 5D) and EARP + GARP (TSSC1) in the steady-state localization of VAMP3 and Syntaxin 6. In the parental cells, GFP:VAMP3 was mainly localized at the plasma membrane, intracellular puncta, and endosomal tubules (Fig. 6D and E). The deletion of *RAB14*, *VPS50*, or *TSSC1* led to a similar phenotype, in which the proportion of cells with VAMP3-positive endosomal tubules decreased (Fig. 6D and E). Because Syntaxin 6 was isolated from EARP complex immunoprecipitations^90^, we tested whether RAB14, EARP, or TSSC1 knockout would perturb its localization. We colocalized Syntaxin 6 with the CI-MPR receptor and observed that its colocalization was perturbed by RAB14, EARP, and TSSC1 KO (Fig. 6F and G). Together, these results functionally link RAB14 to the EARP complex and highlight the extensive role of RAB14 in endosomal trafficking as a regulator of membrane tethering mediated by the EARP complex.

**Figure 6:**
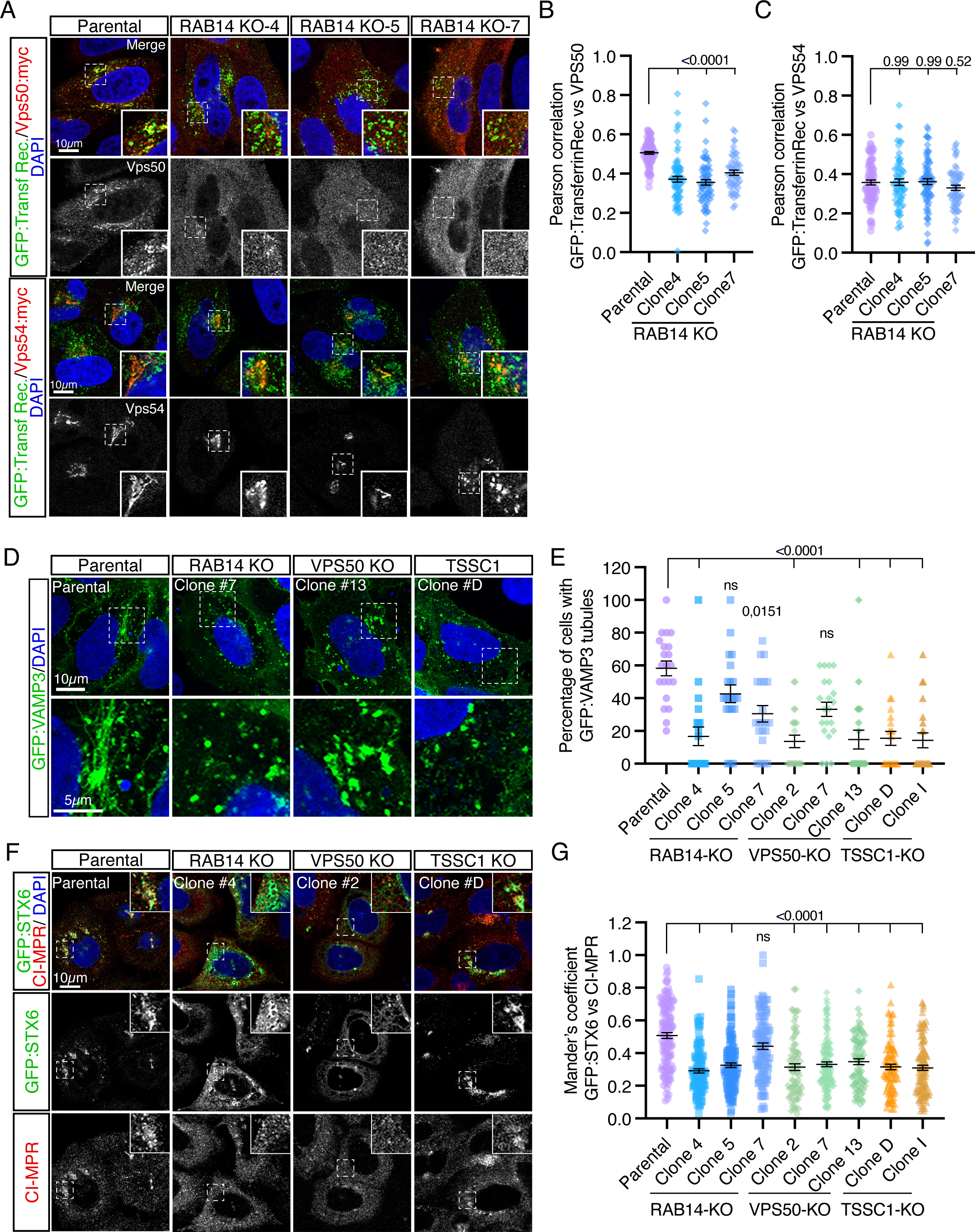
RAB14 is required for EARP-complex endosomal recruitment. **(A)** VPS50 is mislocalized in RAB14 KO HeLa clones. Transiently expressing GFP:Transferrin receptors and (A) VPS50:myc or VPS54:myc HeLa parental and *RAB14* KO cells were fixed and stained. Boxed regions are magnified, and single channels are depicted. Scale bars: 10µm. **(B and C)** Pearson’s correlation between RAB25 and (B) VPS50 or (C) VPS54. Error bars are SEM, individual points represent single cells, *n*=3 independent experiments. **(D)** *RAB14*, *VPS50,* and *TSSC1* KOs modify GFP:VAMP3 localization pattern. Transiently expressing GFP:VAMP3 HeLa parental and knockout cells were fixed and stained. **(E)** Percentage of cells with GFP:VAMP3 tubules in parental or knockout cells. Error bars are SEM, individual points represent single cells, *n*=3 independent experiments. **(F)** *RAB14*, *VPS50,* and *TSSC1* KOs modify GFP:Syntaxin 6 localization pattern. Transiently expressing GFP:Syntaxin 6 HeLa parental and knockout cells were fixed and stained for endogenous CI-MPR. **(G)** Mander’s correlation between GFP:Syntaxin 6 and CI-MPR. Error bars are SEM, individual points represent single cells, *n*=4 independent experiments. Stats used in B, C, E and G were Kruskal-Wallis followed by Dunn’s post test to compare KO lines to the parental line.

### Analysis of most enriched RAB14 interactors identifies a role for RAB14 in regulating SHIP164 and its ortholog UHRF1PB1 localization

Finally, to assess whether single interacting proteins could be identified with confidence from the APEX2 dataset, we plotted the 15 most specific and abundant RAB14 interactors. This highlights known RAB14 interactors along with novel potential binding proteins. The abundant recovery of SHIP164/UHRF1BP1L makes this protein a promising candidate (Fig. 7A). SHIP164 also has a close paralog, UHRFIBP1, and both proteins bind to Vps51^94–96^ directly. Therefore, we tested the interactions between RAB14, SHIP164, and UHRF1PB1. Interestingly, RAB14 interacted with both paralogs (Fig. 7B). Since SHIP164 was expressed less efficiently than UHRF1BP1, we could not determine the binding preference between SHIP164 and UHRF1BP1 for RAB14 (Fig. 7B). While our studies were ongoing, an interaction between RAB14 and SHIP164 was reported^97^. Given our co-immunoprecipitation analysis results and the strong interaction between RAB14 and UHRF1BP1, we examined the relationship between RAB14 and UHRF1BP1. Colocalization analysis demonstrated a high degree of colocalization between activated and wild-type RAB14 and UHRF1BP1 in punctate structures (Fig. 7C and D). This result was consistent with the co-immunoprecipitation analysis data. Based on these results, we tested whether RAB14, EARP, or TSSC1 KO impaired UHRF1BP1 localization. UHRF1BP1 was previously shown to be an endolysosomal^96^ protein, and we detected some degree of co-localization with LAMP2 (Fig. 7E and Suppl. Fig. 5E). This colocalization was affected by RAB14, EARP, and TSSC1 knockout to different degrees between clones (Suppl. Fig.5E). Through these experiments, we noticed that UHRF1BP1 overexpression affected trans-Golgi organization, but we detected good colocalization with punctate TGN46 structures (Fig. 7E and F). This colocalization decreased upon the loss of RAB14, EARP, and TSSC1 (Fig. 7E and F). Based on these results, we conclude that RAB14 interacts with and modulates UHRF1BP1 localization through a probable EARP-dependent mechanism.

**Figure 7:**
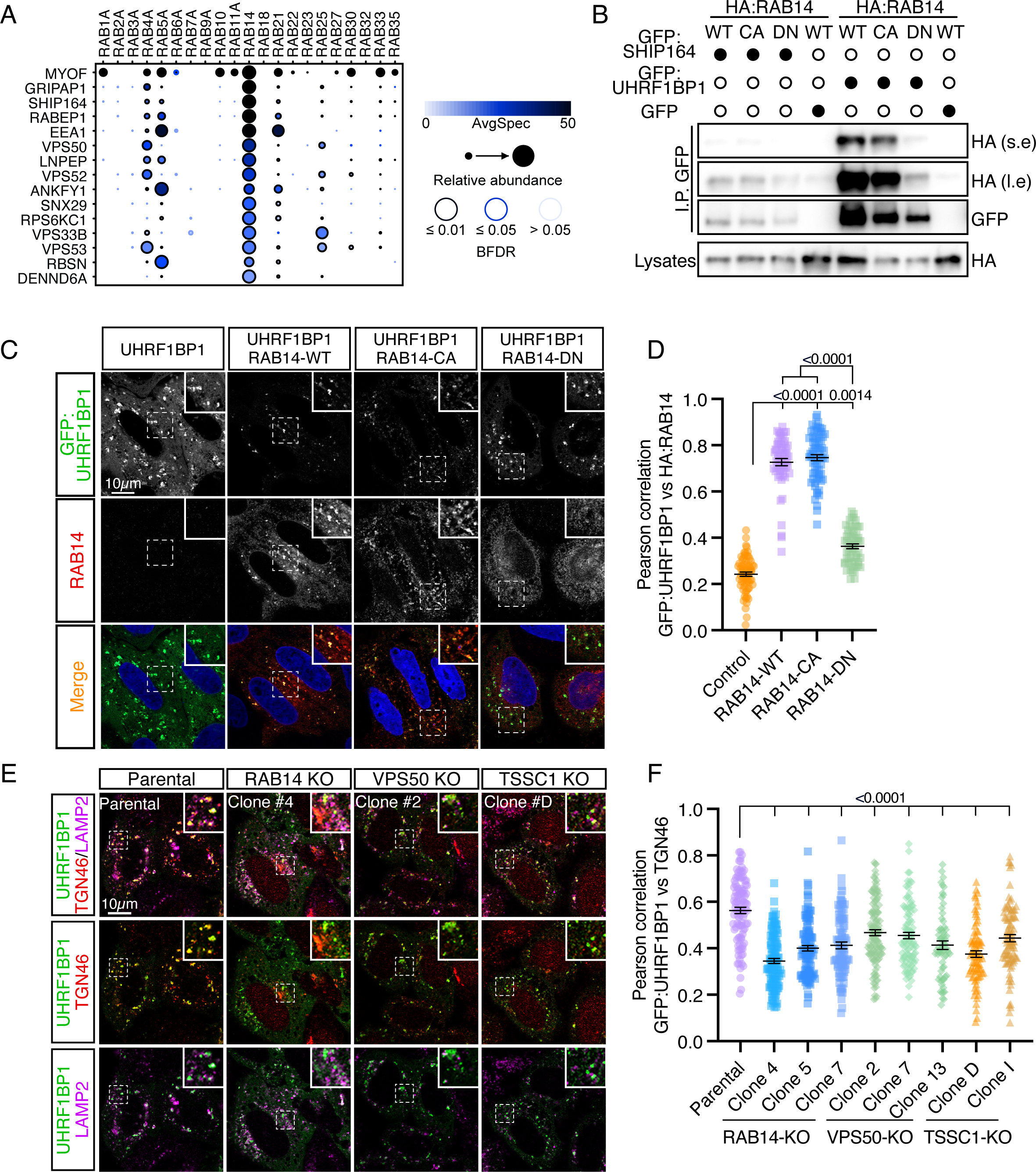
RAB14 interacts with SHIP164 and its paralog UHRF1BP1 and modulates its localization. **(A)** ProHits-viz generated dot plot of the 15 most specific and enriched RAB14 proximal proteins. **(B)** RAB14 interacts with SHIP164 and its paralog UHRF1BP1. GFP-Trap IP of GFP, GFP:SHIP164, and GFP:UHRFBP1 with HA:RAB14 WT, Q70L (CA), and S25N (DN) variants in HeLa cells followed by GFP and HA immunoblots, *n=3* independent experiments. **(C)** RAB14 highly colocalizes with UHRF1BP1. Transiently expressing HA:RAB14 WT, Q70L (CA), and S25N (DN) variants and GFP:UHRF1BP1 HeLa cells were fixed and stained. Boxed regions are magnified, and single channels are depicted. Scale bars: 10µm. **(D)** Pearson’s correlation between RAB14 and UHRF1BP1. Error bars are SEM, individual points represent single cells, *n*=3 independent experiments. **(E)** *RAB14*, *VPS50,* and *TSSC1* KOs modify GFP:UHRF1BP1 localization pattern. Transiently expressing GFP:UHRF1BP1 HeLa parental and knockout cells were fixed and stained for endogenous TGN46 and LAMP2. **(G)** Pearson’s correlation between GFP:UHRF1BP1 and TGN46. Error bars are SEM, individual points represent single cells, *n*=3 independent experiments. Stats used D and F were Kruskal-Wallis tests followed by Dunn’s post test and comparing all groups together (D) or only to the parental line (F).

## DISCUSSION

Proximity labeling has become a powerful tool for robust identification of protein interactomes within a cell^47^. Recent studies have cataloged the interactomes of small GTPases of the Rho^53^ and Arf^54,55^ families. Performing proximity labeling on most members of a protein family allows for the efficient identification of a wide range of novel, specific, and shared interactors, resulting in the discovery of new functions for these GTPases^53–55^. We have previously demonstrated the efficacy of APEX2 labeling for RAB GTPases^49^. In this study, we built an extensive neighboring interaction map of RAB GTPases. Our efforts identified several new interactions and functional relationships.

Using the range of RAB GTPases tested, we observed substantial differences in the number of neighboring proteins between RABs. This may have resulted from the differential accumulation of biotin phenol in the diverse membrane compartments tested. Biotin phenol is hydrophobic; consequently, it would be worth testing alternative substrates that have recently been developed^46^ to label RABs associated with specific compartments, such as endolysosomes. Nonetheless, most RABs, with a few significantly enriched proteins, still displayed proximity with known proximal proteins and were associated with the relevant GO terms, except for RAB11, 22, 23, and 27. The reason for the recovery of RNA-modifying enzymes is unclear, but could be linked to inappropriate localization or membrane insertion. The expression levels of BioID baits have recently been shown to affect the specificity of extracted proteins^98^. Although we used an inducible system that expressed low levels of APEX2:RABs, it is plausible that these four RABs were more sensitive to expression levels, resulting in non-specific localization and labeling of nuclear proteins.

A few studies have cataloged RAB GTPase-interacting proteins^33,35,36,50^, and we directly compared overlapping RABs across different studies. Our dataset intersected to the highest degree with another BioID study in which RAB GTPases were forced to localize at the mitochondria to identify specific binding proteins^50^. Our study and that of Gillingham et al. (2019) identified GO terms linked to membrane trafficking in greater numbers and significance, suggesting that proximity labeling yields more specific interactomes than affinity purification methodologies. We inferred APEX2:RAB localization by comparing our highly significant hits to a BirA-generated cell map^47^. This emphasizes that although different BioID enzymes were used, both datasets were sufficiently robust for most RABs to predict their localization from their associated neighboring proteins.

We capitalized on our extensive dataset and used different metrics to select candidates for validation and functional studies. By comparing hits across all baits with a specificity score (WD score) and an abundance metric, we pursued multiple candidates. Our validation study demonstrated that DENND6A modulates RAB25 localization and that this interaction favors GTP-loaded RAB25. We also showed that DENND6A was required for the RAB25-mediated potentiation of cell migration. Notably, we did not observe a drastic defect in migration of DENND6A KD cells, as observed by Linford et al observed^72^, potentially because, unlike A549 cells, MDA-MB-231 cells do not express N-cadherin. We tested whether RAB25 affects RAB14 colocalization with DENND6A. However, we did not observe a striking difference in RAB14 localization in RAB25 knockout cells compared to parental, suggesting distinct functions to both proteins’ interaction with DENND6A. DENND6A is found both at the recycling endosomes and at the lysosomes^77,82^. While DENND6A is a known GEF for RAB14 at the recycling endosomes^72^, it is quite possible that it could be responsible for RAB25 recruitment to the lysosome. Lysosomal-localized RAB25 was found responsible for the trafficking of active integrins and for their subsequent CLIC3-dependent recycling at the cell’s rear where they promoted cancer cell migration^77^. This is consistent with the reduced cell migration we observed after DENND6A depletion in MDA-MB-231 expressing RAB25, but not in RAB25 negative parental cells. Neither we nor Linford *et al.*^72^, saw a difference in total β1 integrin protein levels in DENND6A-depleted cells. We reasoned that without DENND6A to recruit RAB25, and consequently route β1 integrin to the lysosome, the internalized integrins take a different recycling path that prevents their degradation but does not favor cell migration. These results present exciting hypotheses that need to be tested in future studies.

To further demonstrate the value of the APEX2:RAB-generated dataset, we assessed the proximity of trafficking complexes to RAB GTPases. Again, by assessing the relative abundance and prey specificity, we identified preferential proximity between the EARP complex and RAB14 compared to RAB4. We also used genetic data from the DepMap^92^ consortium to define the potential functional links between these proteins. We validated our MS data through immunoprecipitation and PLA assay results and showed that RAB14 modulates EARP recruitment, affecting the localization of two SNARES previously associated with the EARP complex^90^. These results are consistent with previously observed genetic interactions in *C. elegans*^93^. As RAB14 was recently discovered to regulate trafficking of cationic substances^99^, it suggests a potential role for the EARP complex in this trafficking pathway. Finally, by ranking RAB14’s most abundant and specific proteins, we demonstrated their proximity to the lipid carrier protein SHIP164^97^. We extended these findings to show that the SHIP164 close paralog, UHRF1BP1, also interacted with RAB14, and that its localization was affected by RAB14 and the EARP complex. This opens up the possibility that RAB14 coordinates with the EARP complex for lipid transfer between various types of endosomal populations or microdomains through its capacity to interact with SHIP164 and its paralog UHRF1BP1. The lack of recovery of UHRF1BP1 in the APEX2:RAB14 dataset could be due to lower expression levels of UHRF1BP1 in HeLa cells than SHIP164^92^. Alternatively, RAB14 has been shown to interact indirectly with SHIP164^97^. This is most likely the case for the interaction between RAB14 and UHRF1BP1, which could involve different ‘bridging’ proteins, that would result in an increased distance between RAB14 and UHRF1BP1, leading to decreased labeling.

Our study had some limitations that need to be contextualized. Similar to most proximity labeling studies, we relied on ectopically expressed APEX2-tagged RABs that were adequately localized to their expected cellular compartments. However, we did not assess whether the fusion proteins could rescue knockout cells. Since GFP-tagged RABs are frequently used to rescue RAB knockouts^100^ and APEX2 has a similar molecular weight, APEX2:RAB are likely to be functional; however, we did not demonstrate this. We relied on APEX2-mediated proximity labeling^43^, which requires a short H_2_O_2_ treatment; thus, some RAB interactors may have been missed or gained because of the oxidative stress caused by this treatment. Newer BioID enzymes have been developed, such as TurboID, miniTurbo, UltraID, and AirID, and these could be beneficial in future work, as they do not require H_2_O_2_ treatment or biotin phenol addition^38^. This may lead to a more uniform recovery of preys with various RABs. The MS and bioinformatics pipelines used in our study relied on MS/MS counts and statistical analysis using SAINT^57^. While this has proven valuable for our study^49^ and many others, the democratization of label-free quantitative data-independent MS will most likely improve the specificity of proximity labeling approaches by allowing quantitative measurement of prey levels^101^. This will allow for a better assessment of the binding differences between RABs. Nevertheless, we highlighted and demonstrated numerous interactions by combining different readouts (specificity and abundance) or integrating other databases, such as genetic interactions on DepMap. We hope that by integrating the current dataset with structural predictions, with Alphafold^102^ for example, or with co-expression or co-evolution databases, it will be possible to further refine our proteomic dataset. Despite these limitations, our dataset featured numerous known interactions among RABs, their GEFs, GAPs, and effectors. We validated several novel interactions and identified new functional relationships between the unknown RAB-binding protein pairs. We believe that the proposed RAB proximity map is a valuable resource for the community.

## MATERIAL AND METHODS

### Cell culture

HeLa, HCT116 Flp-in/T-Rex, MDA-MB-231, and MCF7 cells were cultured in Dulbecco’s modified Eagle’s medium (DMEM, 319-005-Cl; Wisent) supplemented with 10% fetal bovine serum (SH30088.03HI; Cytiva) at 37 °C and 5% CO_2_. APEX2 controls and APEX2:RABs were generated by transfecting individual APEX2:RABs with pOG44 plasmid at a ratio of 1:10. Specifically, 900 ng of APEX2:RABs in the pGLAP vector was co-transfected with 100 ng of pOG44 and JetPRIME (101000046; Polyplus), following the manufacturer’s instructions, in individual wells of a 6-well plate. The following day, cells were selected using 150 µg/ml of hygromycin and 5 µg/ml of blasticidin. The selected cells were amplified to yield polyclonal populations used for all individual experiments. All APEX2:RABs proximity labeling was performed in HeLa cells, except for RAB25, which was performed in HCT116 cells because they express RAB25, whereas HeLa cells do not. Cytosolic APEX2 was used in HeLa and HCT116 cells as negative control^49^.

### APEX2:RAB proximity labeling

8.5 × 10^6^ HeLa cells or 10 × 10^6^ HCT116 cells were seeded in 150 mm plates. APEX2:RABs expression was induced on the following day using 10 ng/ml doxycycline for 16 h. After the induction period, DMEM was replaced with full DMEM containing freshly solubilized biotin-phenol (A8011-100(AP); ApexBio) at a final concentration of 500 µM and incubated for 30 min at 37 °C. APEX2 activity was induced by the addition of 2 mM H_2_O_2_ for precisely 1 min. The biotinylation reaction was terminated by transferring cells to ice and performing five washes for one minute with ice-cold quencher buffer (10 mM sodium azide, 10 mM sodium ascorbate, and 5 mM Trolox (53188-07-1; Sigma-Aldrich)). Cells were lysed in 1 mL of cold IP buffer (25 mM Tris-HCl pH 7.4, 1 mM EDTA, 0.1 mM EGTA, 15 mM MgCl_2_, 150 mM NaCl, 2 mM Na_3_VO_4_, 10% glycerol, 1% IGEPAL CA-630, protease inhibitors (P8340; Sigma-Aldrich)), scrapped and left on ice for 20 min, followed by centrifugation at 13,200 RPM for 10 min at 4 °C. Supernatants were collected, and 900 µL of supernatant was incubated with 25 µL of streptavidin Sepharose high performance (17-5113-01; Cytivia) for 2.5 h on a rotating wheel at 4 °C. Following incubation, the beads were washed thrice with complete IP buffer, once with IP buffer lacking IGEPAL CA-630, and five times with 20 mM NH_4_CO_3_. The beads were flash-frozen in liquid nitrogen prior to on-bead digestion for MS. Construct expression and protein biotinylation were validated by western blotting or immunofluorescence.

### Mass spectrometry preparation and analysis

Mass spectrometry analysis was performed as described previously^49^. Briefly, and as mentioned in^49^, the proteins were reduced for 30 min using 10 mM DTT, followed by alkylation for 1 h with 15 mM iodoacetamide (I6125; Sigma-Aldrich). The iodoacetamide was then quenched with 15 mM DTT, and proteins were digested with 1 µg of Pierce MS-grade trypsin (90057; Thermo Fisher Scientific) and incubated overnight at 37 °C with agitation. The digestion was stopped with 1% formic acid, and the supernatant was collected. Residual peptides were eluted with 60% acetonitrile and 0.1% formic acid. The samples were dried, resuspended in 0.1% trifluoroacetic acid (TFA), and desalted using a ZipTip. Ten µl of the sample (2 µg of peptides) resuspended in 1% formic acid were loaded onto a trap column and separated by a PepMap C18 nano column using a linear gradient of 5-25% solvent B (90 % acetonitrile with 0.1% formic acid) over a 4-hour gradient with a constant 200 nl/min flow. Full-scan MS survey spectra were acquired in the profile mode on a QExactive OrbiTrap mass spectrometer with a resolution of 70,000 × using 1,000,000 ions. All unassigned charge states and singly, seven, and eight charged species of the precursor ions were rejected. The lock mass option was used to improve the mass accuracy of the survey scans, and data acquisition was performed using Xcalibur version 2.2 SP1.48.

After sample analysis by nano LC-MS/MS, the peptides and proteins were identified using MaxQuant version 1.5.2.8, and the Uniprot database (Homo sapiens, 16/07/2013, 88354 entries). Trypsin/P was set as the enzyme, with cleavages allowed on arginine or lysine preceding proline and a maximum of two miscleavages. Mass tolerances of 7 and 20 ppm were used for the precursor and fragment ions, respectively. The modifications included fixed carbamidomethyl on cysteine, variable oxidation of methionine, and N-terminal acetylation. All proteins had to pass a False Discovery Rate (FDR) of less than 1% to ensure reliable identification. Any proteins whose hits in the forward database were not 100-fold higher than those in the reverse database were excluded. Only proteins with two or more peptides were analyzed using SAINT and frequent contaminants were removed. Three independent biological repeats per RAB were performed for each RAB. Correlations between repeats were established, and the level of correlation varied significantly depending on the RAB tested (Extended Table 1). The correlation map between repeats and the different RAB was established by comparing all captured preys with a SC ≥ 0.75. and by calculating Jaccard indexes for all combinations and a Spearman’s rank correlation for repeats of an individual RAB. Hence, we excluded poorly correlated replicates and retained a minimum of two replicates in the final analysis. To identify high-probability neighboring proteins, SAINTExpress was first run on the Crapome platform using APEX2 as the user control, and then compared to each APEX2:RAB. Only proteins with SAINT scores equal to or above 0.95 were designated high-probability protein neighbors (Extended Table 1). Raw SAINT results were used to create dot plots or generate WD scores on the Prohits-Viz platform. We utilized the two most correlated experiments for the WD score calculations, given that the number of repeats is a factor in the score calculation. Finally, to account for experimental variations and different labeling potencies of the different RABs, all samples were normalized to the total number value on Prohits-Viz for dot plot representations. The complete and unfiltered dataset is available in the PRIDE database with the identification number PXD057404.

### Generation of DNA constructs

Human RABs were PCR amplified with the Phusion Hot Start Flex 2X master mix (M0536; New England Biolabs) from a cDNA library generated from mixed total RNA extracted with Trizol reagent (15596026: Thermo Scientific) from HeLa, HCT116, and MCF7 cells. PCR-amplified RABs were then cloned in-frame with MYC:APEX2 through Gibson ligation using the In-Fusion HD kit (639650; Takara Bio) in a modified pGLAP1 vector^49^ to yield the MYC:APEX2:RABs expression vector. EGFP:RAB32WT was generated by subcloning RAB32 from the MYC:APEX2:RAB32 plasmid and transferring it to pEGFP-C1 by Gibson ligation. GFP:RAB32CA (Q85L) and GFP:RAB32DN (T39N) were generated by site-directed mutagenesis using the Q5 site-directed mutagenesis kit (E0554; New England Biolabs). ALS2 with a single C-terminal HA tag was synthesized by Twist Biosciences and inserted into the NotI/NheI sites of the pTwist CMVPuro plasmid. 3xHA:GRAB was also synthesized and inserted into the pTwist CMVPuro plasmid with an N-terminal 3xHA tag. EGFP:DENND6A was PCR-amplified from the pooled cDNA and inserted into the pEGFP-C1 plasmid via Gibson ligation. Different expression plasmids were generated for hRAB25. Transient expression was performed using RAB25 inserted into the pCDNA3-Nterm 3xHA plasmid. RAB25 Q71L was generated using a Q5 site-directed mutagenesis kit and further modified to yield the T26NQ71L double mutant RAB25. RAB25 was inserted into the pLVX-GFP lentiviral vector for stable cell expression to yield a pEGFP:RAB25 fusion protein. GFP:RAB14 was generated via PCR amplification of RAB14 from the MYC:APEX2:RAB14 plasmid and subcloned into the pEGFP-C1 plasmid. CA (Q70L) and DN (S25N) cells were generated as described previously. EGFP-Rab14WT, CA, and DN were also subcloned into pCDNA3 Nterm 3xHA by PCR amplification and Gibson ligation. GFP:Syntaxin 6 was PCR-amplified from HeLa cDNA and inserted into the pEGFP-C1 plasmid via Gibson ligation. COG1, COG4, and COG6 were PCR-amplified from a mixed cDNA pool and cloned into the pCDNA3-3xHA plasmid using Gibson ligation. gRNA cloning was performed by digesting pX330A-1x2 (58766; Addgene) with Bbs1 (R3539; New England Biolabs). Oligos with optimal gRNA sequences for RAB14, VPS50, TSSC1, and RAB25 obtained from Brunello table^103^, were annealed and ligated with T4 DNA ligase (M0202; New England Biolabs). All plasmids generated in this study were sequence-validated. The following plasmids were obtained from Addgene: HA:RABIN8 (118071), GFP:RAB2A (49541), VPS50-13xMYC (175093), VPS54-13xMYC (175095), pTfR-mNeonGreen (129608), and GFP:VAMP3 (42310). Pietro de Camilli and Karin Reinisch kindly provided GFP:SHIP164 and GFP:UHRFPB1. Juan Bonifacino kindly provided V5:VPS53.

### Immunoprecipitations and immunoblots

APEX2:RAB expression and the ability to biotinylate neighboring proteins were evaluated by seeding 100,000 cells per well in a six-well plate. The following day, cells were incubated in biotin-phenol, stimulated with or without H_2_O_2_, and quenched as described above. Cells were then lysed in RIPA buffer (10 mM Tris-HCl pH8, 1mM EDTA, 0.5 mM EGTA, 1% Triton X-100, 0.1% Sodium Deoxycholate, 0.1% SDS, 140 mM NaCl and 1x protease inhibitor cocktail) for 20 min on ice. Cell lysates were spun at 16,000 *g* for 10 min at 4 °C. Supernatants were collected and protein concentrations were determined using the Pierce BCA assay (23225; Invitrogen (Thermo Fisher Scientific)). Equal amounts of proteins were loaded onto SDS-PAGE gels (range between 8-13% based on the molecular weight of proteins analyzed) and transferred onto PVDF membranes (10600021; Cytiva) using a TurboBlot (1704150; Bio-Rad) semi-dry transfer apparatus.

For coimmunoprecipitations of transfected proteins, 80,000 cells per well (HeLa) and 300,000 cells (MCF7) were seeded in a six-well plate. The following day, cells were transfected with 1 ug of DNA (total) using JetPrime reagents, according to the manufacturer’s instructions. The next day, cells were processed as in^104^. Briefly, cells in full medium were lightly fixed in 0.5% formaldehyde (final concentration) for 15 min with rocking at room temperature. Formaldehyde was quenched with 125 mM glycine (final concentration) at pH 3 for 5 min at room temperature. Cells were washed with ice-cold PBS twice and lysed with 300 µL of IP buffer (25mM Tris-HCl pH 7.4, 150mM NaCl, 1mM EDTA, 5mM MgCl_2_, 5% glycerol, 1% Igepal CA-630 and protease inhibitors) for 20 min on ice. Cells were scraped and spun at 16,000 *g* at 4 °C for 10 min, and supernatants were collected and quantified. Equal amounts of cell lysates were processed for IPs and incubated with glutathione Sepharose 4B beads (17-0756-01; Cytiva) for 1h at 4 °C under rotary agitation to preclear the lysates. Glutathione sepharose beads were removed by centrifugation, and cell lysates were incubated for 1 h at 4 °C under rotary agitation with GFP Trap beads pre-blocked overnight with lysates from non-transfected cells. The beads were washed thrice with CO-IP lysis buffer and once with a buffer without NP-40. For HA-IPs, 2 µg of anti-HA tag antibody (sc-7392; Santa Cruz Biotechnology) was added to cell lysates and incubated for 1h at 4 °C with rotation, followed by the addition of 25 µL of equilibrated protein A sepharose (ab193256; Abcam). The incubation was continued for another 1 h at 4 °C with rotation. IPs were washed 4x with ice-cold IP buffer. For immunoprecipitation of endogenous proteins, 30 µL of equilibrated A/G agarose beads (sc-2003; Santa Cruz Biotechnology) was mixed with 1 ug of anti-VPS50 or with 1 ug of total goat antibodies and incubated overnight at 4°C with rotation. HeLa cells lysates were extracted from 80 to 90% confluent 100 mm petri dishes. Briefly, cells were washed twice with cold PBS and lysed in 500 µL of HEPES lysis buffer (HEPES 20 mM, NaCl 150 mM, EDTA 1 mM, Triton-x-100 1%, protease inhibitors 1X). Protein collection was performed as mentioned above. Antibody-bound beads were washed thrice in HEPES lysis buffer and incubated with 500 µL of cell lysates overnight at 4°C with rotation. Beads were washed twice with HEPES lysis buffer and again twice with 20 mM HEPES. For all IPs, beads were resuspended in 2x Laemmli SDS-loading buffer, heated for 5 min at 95 °C and processed for immunoblot.

The PVDF membranes were processed for immunoblot analysis according to the general Cell Signaling Technology (CST) western blotting protocol. The antibodies used were anti-MYC (1:1000, 71D10), anti-GAPDH (1:1000, D16H11), anti-HA (1:1000; C29F4), anti-VPS53 (1:2000, 2B12C3), and anti-RAB25 (1:1000, D4P6P; Cell Signaling Technology). Anti-GFP (1:1000, SC-9996) was purchased from Santa Cruz Biotechnology, anti-V5 (1:2000; V8012), anti-tubulin (1:5000; T9026), and anti-RAB14 (1:500; HPA026419l) were purchased from Sigma-Aldrich and anti-VPS50 (1:1000; A303-324A-T) was purchased from Bethyl Laboratories. Horseradish peroxidase (HRP)-conjugated Streptavidin (1:100; N100) was purchased from Thermo Fisher Scientific, and secondary antibodies against mouse (1:10 000; 115-035-146) and rabbit (1:10 000; 111-035-144) were purchased from Jackson ImmunoResearch. Mouse Anti-rabbit IgG conformation-specific (1:2000, L27A9; Cell Signaling Technology) was used to label proteins close to 25 and 50 kDa that were immunoprecipitated with a rabbit antibody. Protein detection by chemiluminescence was performed using Luminata Forte (ELLUF0100; Millipore) or Clarity Max (1705062; Bio-Rad) substrates, and images were acquired on an Azure C280 Chemidoc.

### Immunofluorescences

For APEX2:RAB IFs, 75 000 Flp-in/T-Rex cells were seeded on glass coverslips (72230-01; Electron Microscopy Sciences), and APEX2:RAB expression was induced for 16 h, as described above. Immunofluorescence was performed according to the general immunofluorescence protocol provided by Cell Signaling Technology. Briefly, cells were washed twice with PBS and incubated in 4% paraformaldehyde (15713; Electron Microscopy Science) in PBS for 15 min at room temperature, followed by three 5 min washes in PBS. Cells were permeabilized and blocked for 60 min in blocking buffer (5% goat serum, 0.3% Triton X-100 in PBS). Cells were rinsed once with antibody buffer (1% BSA, 0.3% Triton X-100 in PBS) and incubated overnight at 4 °C with primary antibodies diluted in antibody buffer. The next day, the cells were washed 3 times for 5 min with PBS. The cells were rinsed with antibody buffer and incubated with secondary antibodies in antibody buffer for 2 h at room temperature, followed by three 5 min washes in PBS. Cells were mounted on slides in 4 µL of slow fade gold mounting media containing DAPI (S36938; Invitrogen (Thermo Fischer Scientific)). Slides were kept at 4 °C in the dark before image acquisitions.

30,000, 100,000, and 75,000 Hela, MCF7, and MDA-MB-231 cells, respectively, were plated as described above. If required, the cells were transfected with JetPrime according to the manufacturer’s instructions. DNA quantity was adjusted based on the JetPrime cell line database. Cells were fixed and immunofluorescence analysis was performed as described above. Antibodies used throughout the study were anti-MYC (1:500; 71D10; Cell Signaling Technology), anti-transferrin receptor (1:100; D59X; Cell Signaling Technology), anti-EEA1 (1:500; C45B10; Cell Signaling Technology), anti-HA (1:500; C29F4; Cell Signaling Technology), anti-TGN46 (1:100; Clone 2F7.1; Genetex), anti-CIMPR (1:100; MCA2048; BioRad), anti-LAMP1 (1:15; 1D4B; DHSB), anti-KDEL (1:200; EPR12668; Abcam), anti-TOM20 (1:200; 612278; BD Biosciences), anti-total ≥1-integrin (1:100; P5D2; DSHB), anti-active ≥1-integrin (1:100; clone 12G10; Millipore Sigma). F-actin was labelled using Alexa-Fluor-conjugated phalloidin purchased from Invitrogen (1:250; A22283; Thermo Fisher Scientific). Secondary antibodies (1:250) were purchased from Invitrogen (Thermo Fisher Scientific) and conjugated with Alexa Fluor 488, 546, and 647 (A11034, A11029, A11035, A11030, A21236, and A21245, respectively).

Total and active β1 integrins in MDA-MB-231 cells were determined similarly to the above protocol with minor adjustments. MDA-MB-231 cells were permeabilized in PBS containing 0,05% saponin and 10% goat serum. Maximum intensity projections of three Z positions (at 1.5 μm from the center on each side) were acquired, and β1 integrin mean intensity per cell was analyzed using CellProfiler software.

### Proximity ligation assay (PLA)

A total of 20 000 HeLa cells were plated on circular coverslips in 24-well plates containing complete medium. The following day, the cells were transfected as described for immunofluorescence analysis with 50 ng of pEGFP-C1 or 500 ng of the indicated plasmids. Cells were fixed with 4% paraformaldehyde for 15 min, washed three times with PBS, and permeabilized with a solution of PBS containing 5% goat serum and 0.3% triton X-100 for 1h at room temperature 30 h post-transfection. Cells were labeled overnight at 4 °C with mouse anti-GFP (1:1500; sc-9996; Santa Cruz Biotechnology), rabbit anti-HA, or mouse anti-MYC (1:100; DHSB) in PBS with 2% BSA (800-095-EG; Wisent) and 0.3% triton X-100. Proximity ligation assay was performed using the Duolink *in situ* PLA kit (DUO92101; Sigma-Aldrich) following the manufacturer’s recommendations. Coverslips were mounted on microscope slides using slowFade gold antifade mountant with DAPI. PLA puncta were counted in GFP+ cells using an unbiased automated pipeline in CellProfiler.

### Cell migration assays

The bottom wells of a 24-well plate were coated with 10 µg/mL of human fibronectin (354008; Corning) for 1h at 37 °C. Wells were rinsed with PBS and then allowed to dry before placing silicone culture inserts (80209; IBIDI) with a 500 µm cell-free gap. 90,000 WT or stably expressing RAB25 MDA-MB-231 cells were seeded into each well and allowed to adhere and grow for 24 h in complete medium. The following day, the silicone inserts were removed, the cells were gently washed with warm PBS, and the culture medium was replaced with DMEM supplemented with 1% FBS to restrict cell proliferation. Cell migration was captured using time-lapse imaging with Cell Discoverer 7 (ZEISS) and analyzed using the FIJI Image J MRI_Wound_Healing Tool plugin to establish the percentage of gap closure.

### CRISPR/Cas9 knockout cell generation, stable cell lines, and siRNA

HeLaM-knockout clones were generated as follows. 100,000 HeLa M cells were plated per well in a 6-well plate and transfected the following day at a ratio of 1:15 with the pEGFP-Puro plasmid (45561; Addgene) and gRNA-containing plasmid (pX330A-1×2). The following gRNA sequences were used: CACAATTGGTGTTGAATT (RAB14), CATTGCAGACAGGTCTTCAA (VPS50), and CCATCAAGCGGGTGAAATCT (TSSC1). The following day, the media was changed to complete DMEM containing 1 µg/mL puromycin, and cells were incubated for 36 h. Resistant cells were detached using trypsin and counted, and single-cell suspensions were added to each well of a 96-well plate. Single-cell clones were amplified and used in subsequent experiments. MCF7 knockout cells for RAB25 were established as described above; however, we used the knockout population instead of clones. The three gRNAs were co-transfected with the pEGFP-puromycin plasmid from Addgene. The following gRNA sequences were used: gRNA1 (GCTTGGTTAGGTCAAACACC), gRNA2 (CACGAGCATGACGACGATCG) and gRNA3 (CGGTGCCCAACATCACAGTG).

Stable MDA-MB-231 cells were generated by lentiviral transduction. Briefly, HEK293T cells in a 15 cm dish were transfected with pMD2G, pCMVdR8.2, and EGFP:RAB25-pLVX plasmids using polyethyleneimine (PEI) (23966; Polysciences). The supernatant was collected and filtered through a 0.45 µM cellulose acetate filter, and lentiviruses were conserved at -80 °C 72 h post-transfection. MDA-MB-231 cells were transduced by incubating 30,000 cells in a 6-well plate with 750 µL of viral supernatant, 750 µL of OptiMEM (11058021; Gibco (Thermo Fisher Scientific)), and 1 µL of polybrene overnight at 37 °C. The following day, cells were selected with 1 µg/mL of puromycin for three days and maintained in completed DMEM containing 0.5 µg/µL of puromycin.

For siRNA-mediated knockdown, 30,000 parental or stably expressing RAB25 MDA-MB-231 cells were cultured in a 6-well plate in complete medium (DMEM, 10% FBS, penicillin/streptomycin) overnight to allow cell adhesion. DENND6A was knocked down using ON-TARGET-PLUS siRNA (J-016505-05-0005 and J-016505-06-0005; Horizon Discovery) at a final concentration of 12.5 nM and transfected with the DharmaFECT 4 reagent (T-2004-02; Horizon Discovery), as per the manufacturer’s instructions. Media containing the siRNA was removed 48 h after the transfection, and cells were allowed to grow in complete media for another 24 h before migration assay or β1 integrins immunofluorescence. We were unable to identify antibodies specific to DENND6 or TSSC1. Thus, siRNA knockdown validation or analysis of knockout efficiencies was performed by RT-qPCR or sequencing of the clonal cell line, respectively.

### qRT-PCR quantifications

RNA from MDA-MB-231 WT and RAB25+ cells was extracted 72 h post-siRNA transfection using GeneJET RNA purification columns (K0732; Invitrogen) following the manufacturer’s instructions. RNA purity and concentration were assessed using a NanoDrop 2000 spectrophotometer (Thermo Fisher Scientific). The Maxima First Strand cDNA Synthesis Kit (K1642; Invitrogen (Thermo Fisher Scientific)) was used according to the manufacturer’s instructions to reverse transcribe 500 ng of RNA into cDNA, including the pre-removal step for genomic DNA. The amplification efficiency and optimal annealing temperature of the primer pairs for *DENND6A* (For 5’ CCAGGCCGTGGAGGTAAT 3,’ Rev 5’

CCCTCCTCCCAGAAGACTGT 3’) and *TBP* (For 5’ GGGGAGCTGTGATGTGAAGT 3,’ REV

5’ GGAGAACAATTCTGGGTTTGA 3’) were evaluated with a template and a temperature standard curve, respectively. qPCR was performed using the Luna Universal qPCR master mix (M3003; New England Biolabs) following the manufacturer’s recommendations. All samples were amplified in duplicates using a Roche <u>L</u>ightCycler 96 thermocycler. *DENND6A* expression was calculated using the 2(-Delta C(T)) method and normalized to the reference gene *TBP*.

### Image analysis and statistics

Images were acquired on a Zeiss LSM880 equipped with a 40x 1.4NA plan Apo objective or a Cell Discoverer 7 using a 20x 0.5NA WD plan-apochromatic objective with the phase gradient adaptive option for wound healing imaging. The settings for fluorescence imaging were set to optimize the pixel size and ensure an adequate distribution of pixel intensities. As such, pixel saturation was minimized, and no offset was added during acquisition, given the good signal-to-noise ratio of the system. Similar imaging settings were utilized within each experimental set to ensure adequate image comparison. The images displayed in the manuscript were first exported as tiff files using Zen Blue, and then processed using Adobe Photoshop. In Photoshop, the images were thresholded and cropped using linear thresholding. All images were processed similarly for the experimental set. The figures were assembled using Adobe Illustrator.

Image analysis was performed using Cell Profiler^105^. Briefly, pipelines were generated to identify single objects (PLA dots, puncta, or cells) depending on the experiment, and Pearson or Manders coefficients were calculated using the colocalization module. Staining intensities were measured with the ‘measure intensity’ module on cells manually identified to ensure appropriate measurements for each cell. The presence of GFP:VAMP3 tubules was manually determined, and the percentage of cells with tubules per field was generated.

All experiments were performed at least three times independently, unless otherwise mentioned. We first assessed the normality of the various datasets using the Shapiro–Wilk test. Statistical tests were adjusted for parametric or non-parametric distributions. Multiple conditions were compared using one-way analysis of variance (ANOVA), followed by Dunnett’s multiple comparison test. Nonparametric Kruskal–Wallis test was adjusted using Dunn’s multiple comparison test. Each time the effect of 2 independent variables was compared on a measured outcome, a 2-way ANOVA followed by Sidak or Tukey’s multiple comparison test was performed. All statistical analyses and graphical representations were performed using Prism 10 software (GraphPad Software). The graph points represent single cells or fields collected from independent experiments.

Metascape analysis was performed by inputting all proteins with a Saint score ≥0.95 using the express analysis option. GO term enrichment was performed similarly using ShinyGo. To infer the APEX2:RAB localization from the HumanCellMap dataset, we directly uploaded the SaintExpress file to the upload and analysis module. Frequent flyers were ignored, and the number of preys analyzed were limited to 150. A heat map was generated by selecting the top four compartments and determining the percentage of prey within these compartments. Additionally, if the localization was specific, a maximum of 17 prey items were used. The heatmap corresponds to the percentage frequency of each prey item in each compartment. Overlap between RAB interactome studies was achieved by selecting overlapping RAB GTPases and combining both GDP- and GTP-identified hits (only for Gillingham et al. (2019)). We selected the best ortholog match from the DIOP Ortholog Finder to convert gene names to human genes because Gillingham et al. (2014) and Li et al. (2016) used fly and mouse cells, respectively. Finally, we applied selectivity factors for each study: SaintScore ≥ 0.95 for this study, S score ≥ 5 for Gillingham et al. (2014), WD score of ≥ 10 Gillingham et al. (2019) and used the high-confidence interactor list from Li et al. (2016). Scripts for these analyses and figures are available at https://github.com/Labo-MAB/RAB_interactomes.

## Acknowledgments

We thank the laboratory members for their assistance throughout the study and Dominique Lévesque for his help with the mass spectrometer. We thank Drs. Juan Bonifacino, Pietro De Camilli, and Karin Reinish for generously providing various plasmids. We used the Université de Sherbrooke microscopy and proteomic cores for imaging and MS. S.J., M.A.B. and F.M.B. are members of the Fonds de Recherche du Québec Santé (FRQS)-funded Centre de Recherche du CHUS and members of the Institut de Recherche sur le Cancer de l’Université de Sherbrooke (IRCUS). S.J. is also a member of the regroupement québécois de recherche sur la fonction, l’ingénierie et les applications des protéines (PROTEO). We would like to thank Editage (www.editage.com) for English language editing. This work was supported by the Centre de Recherche Médicale de l’Université de Sherbrooke institutional research chair and a Discovery Grant from the Natural Science and Engineering Research Council (NSERC - 03730). S.J., M.A.B., and F.M.B. received salary awards from FRQS. F.B. and M.M. are recipients of FRQS and NSERC master scholarships, respectively.

