## Supplemental figures for "A Proximity MAP of RAB GTPases"

### SUPPLEMENTARY FIGURE LEGENDS

**Supplementary Figure 1: APEX2:RAB cell line validation. (A)** Phylogenetic tree representation of RABs tested in this study. The tree representation is from[29]. Green indicates that the tested RAB is expressed and can biotinylate proteins, while red signifies defective expression or biotinylation. **(B)** Most APEX2:RABs can biotinylate endogenous proteins. Total biotinylated proteins revealed by streptavidin western blotting in APEX2:RAB Flp-In/T-REx HeLa cells. **(C)** ProHits-viz generated dot blot of individual RABs self-biotinylation and their respective abundance and statistical significance.

**Supplementary Figure 2: Associated RAB proximal proteins can infer RAB localization and general analyses of the RAB proximity map. (A)** The topmost specific proteins for each RAB were used to identify the potential localization of the tested APEX2:RAB using the cell map dataset[47]. **(B)** Go-terms enrichment analysis of all proteins identified with a SC above 0.95 for all RABs. **(C)** Upset plots of three RAB interactome studies compared to the new APEX2:RAB dataset presented in this study, detailing the number of overlapping GO-terms. **(D and E)** Heat maps of WD scores for (A) GEFs and (B) GAPs identified enriched with a Saint score of $\geq 0.95$ for the various RABs. RABs for which no GEFs or GAPs with a SC $\geq 0.95$ are present were omitted from the heatmaps for representation purposes.

**Supplementary Figure 3: DENND6A and RAB25 are functionally linked. (A)** Transiently expressing GFP:RAB25 T26NQ71L co-transfected with either GFP or GFP:DENND6A MCF7 cells were fixed and stained. Boxed regions are magnified, and single channels are depicted. Scale bar: 10µm, *n=3* independent experiments. **(B)** DENND6A and RAB14 colocalization does not require RAB25. Transiently expressing GFP:DENND6A and HA:RAB14 MCF7 cells were fixed and stained. Boxed regions are magnified. Scale bar: 10μm. **(C)** Pearson's correlation between DENND6A and RAB14. Error bars are SEM, individual points represent single cells, *n*=3 independent experiments. Validation of the cell lines used to study RAB25 **(D)** Low RAB25 expression levels in the RAB25 MCF7 KO cell population. Immunoblot analysis of endogenous RAB25 and GAPDH. **(E)** High GFP:RAB25 expression in the MDA-MB-231 stable cell population. Immunoblot analysis of GFP:RAB25 and GAPDH. **(F)** DENND6A expression is significantly increased in GFP:RAB25-expressing MDA-MB-231 cells. Quantitative RT-PCR 2(-Delta C(T)) analysis of both MDA-MB-231 cell lines. **(G)** DENND6A siRNAs efficiently decrease DENND6A mRNA levels. Quantitative RT-PCR analysis of DENND6A mRNA levels in cells transfected with a scramble siRNA or two independent DENND6A-targeted siRNA. **(H-K)** DENND6A depletion does not extensively affect total or active β1-integrin levels. Parental and stable GFP:RAB25 expressing MDA-MB-231 cells transfected with control, or DENND6A targeted siRNA were fixed and stained for endogenous (H) total β1-integrin or (J) active β1-integrin and F-actin. Mean fluorescence intensity par cell (arbitrary units) of (I) total β1-integrin or (K) active β1-integrin. Error bars are SEM, individual points represent single cells, *n*=3 independent experiments. Stats used in C and F were two tail t-test and G, I and K were 2-way ANOVA with Sidak post test comparing DENND6A siRNA samples to siScramble control's.

**Supplementary Figure 4: Various membrane trafficking regulatory complexes show proximity to various RAB GTPases.** **(A)** Heat maps of WD scores for various membrane-associated trafficking complexes enriched with a Saint score $\geq 0.95$ for the RABs tested. **(B)** PLA immunofluorescence images show the proximities between wild-type GFP:RAB2A and

COG1:HA, COG4:HA, and COG6:HA in HeLa cells. All constructs were also transfected individually and probed with both antibodies. PLA puncta are shown in red and nuclei were stained with DAPI (blue). Dotted lines define individual cells. **(C)** Quantification of the PLA puncta in (B). Individual points represent single cells, and the error bars are the SEM (*n*=3 independent experiments). Control represents the single GFP:RAB2A transfection. Stats used in C were a 2-way ANOVA followed by Tukey post test

**Supplementary Figure 5: RAB14 shows proximity to EARP and GARP complexes by proximity ligation assay. (A)** PLA immunofluorescence images showing the proximities between wild-type HA:RAB14 WT, Q70L (CA), and S25N (DN) variants and VPS50:13xMYC or VPS54:13xMYC in HeLa cells. All constructs were also transfected individually and probed with both antibodies as controls. PLA puncta are shown in red and nuclei were stained with DAPI (blue). Dotted lines define individual cells. **(B)** Quantification of the PLA puncta in (A). Individual points represent single cells, and the error bars are the SEM (*n*=3 independent experiments). Control represents the single GFP:RAB14 WT, CA or DN transfections. **(C-D)** RAB14 and VPS50 expression levels in the (C) RAB14 and (D) VPS50 HeLa KO clones. Immunoblot analysis of endogenous RAB14, VPS50, and TUBULIN. **(E)** Pearson's correlation between GFP:UHRF1BP1 and LAMP2. Error bars are SEM, individual points represent single cells, *n*=3 independent experiments. Stats used in B were 2-way ANOVA with a Tukey post test and in E was Kruskal-Wallis with Dunn's post test.

### Supplementary Figure 1

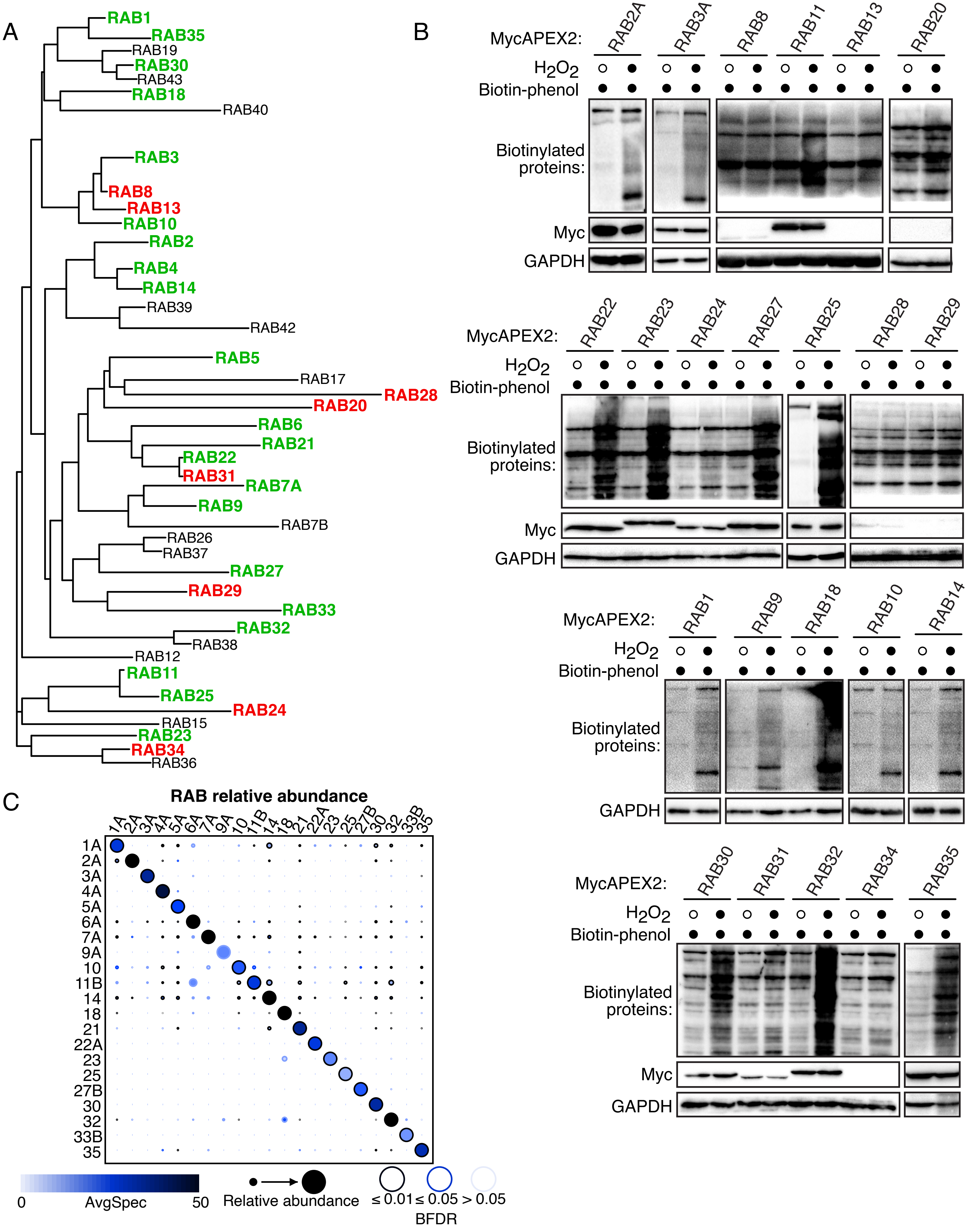

### Supplementary Figure 2

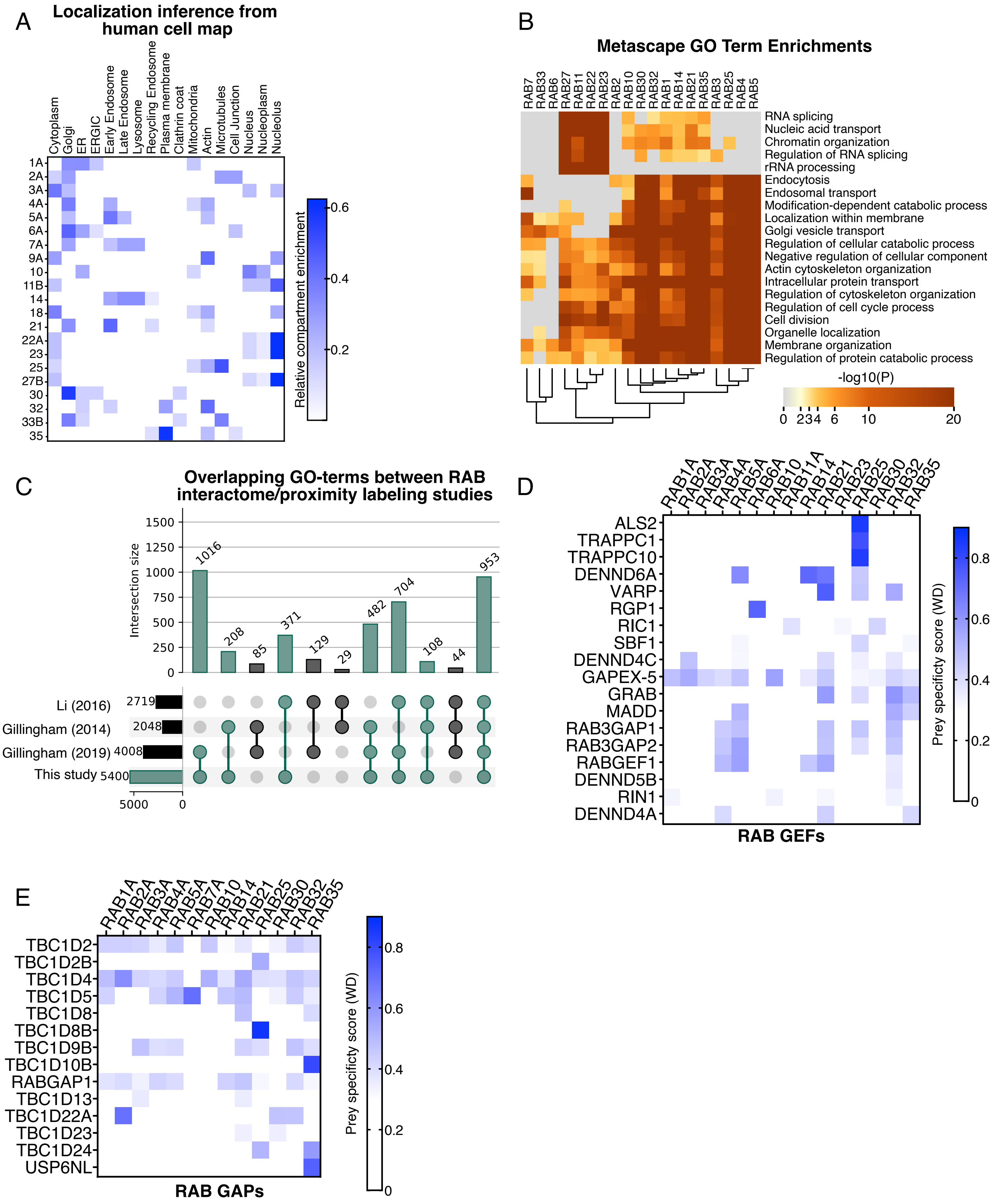

**Supplementary Figure 3**

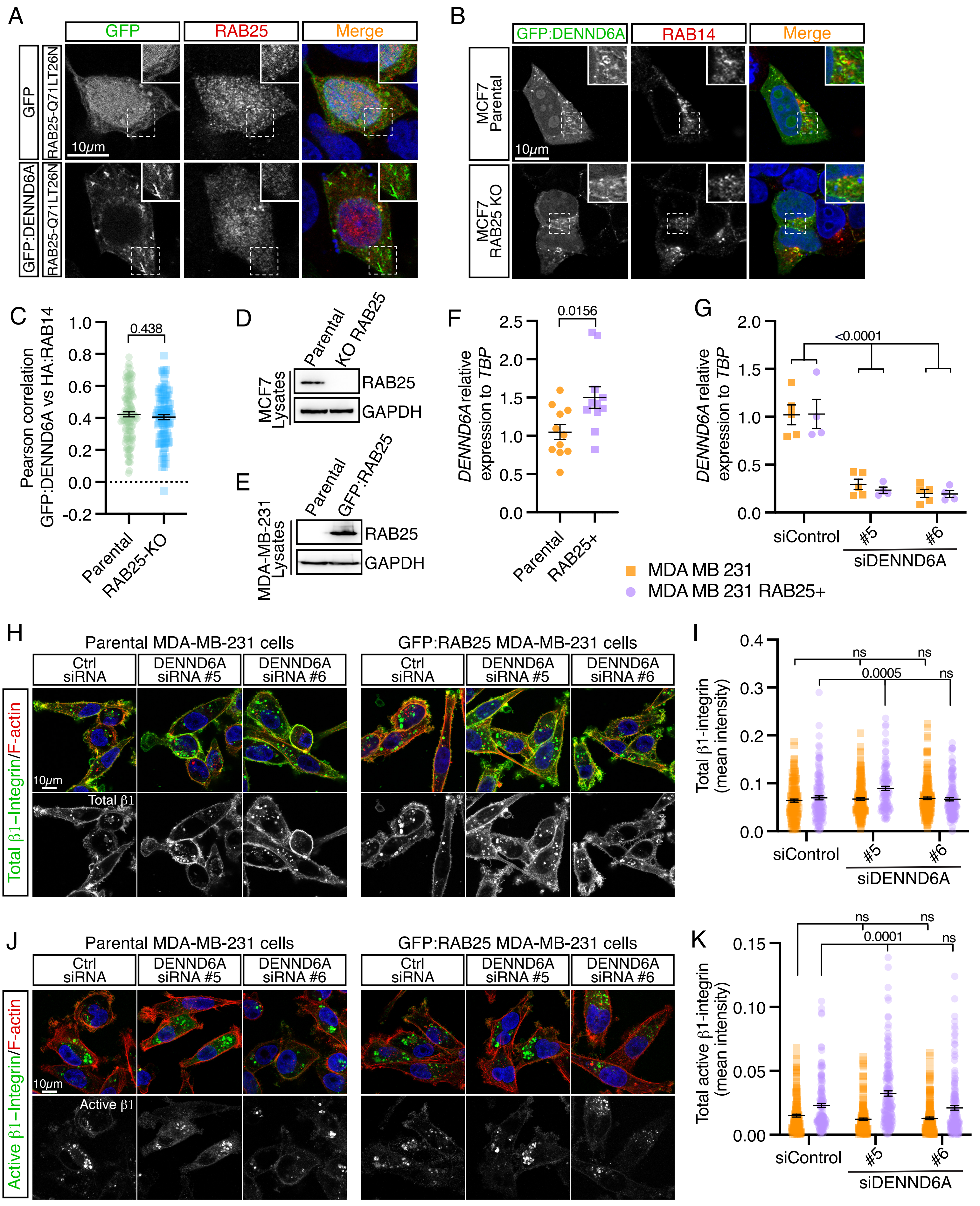

### Supplementary Figure 4

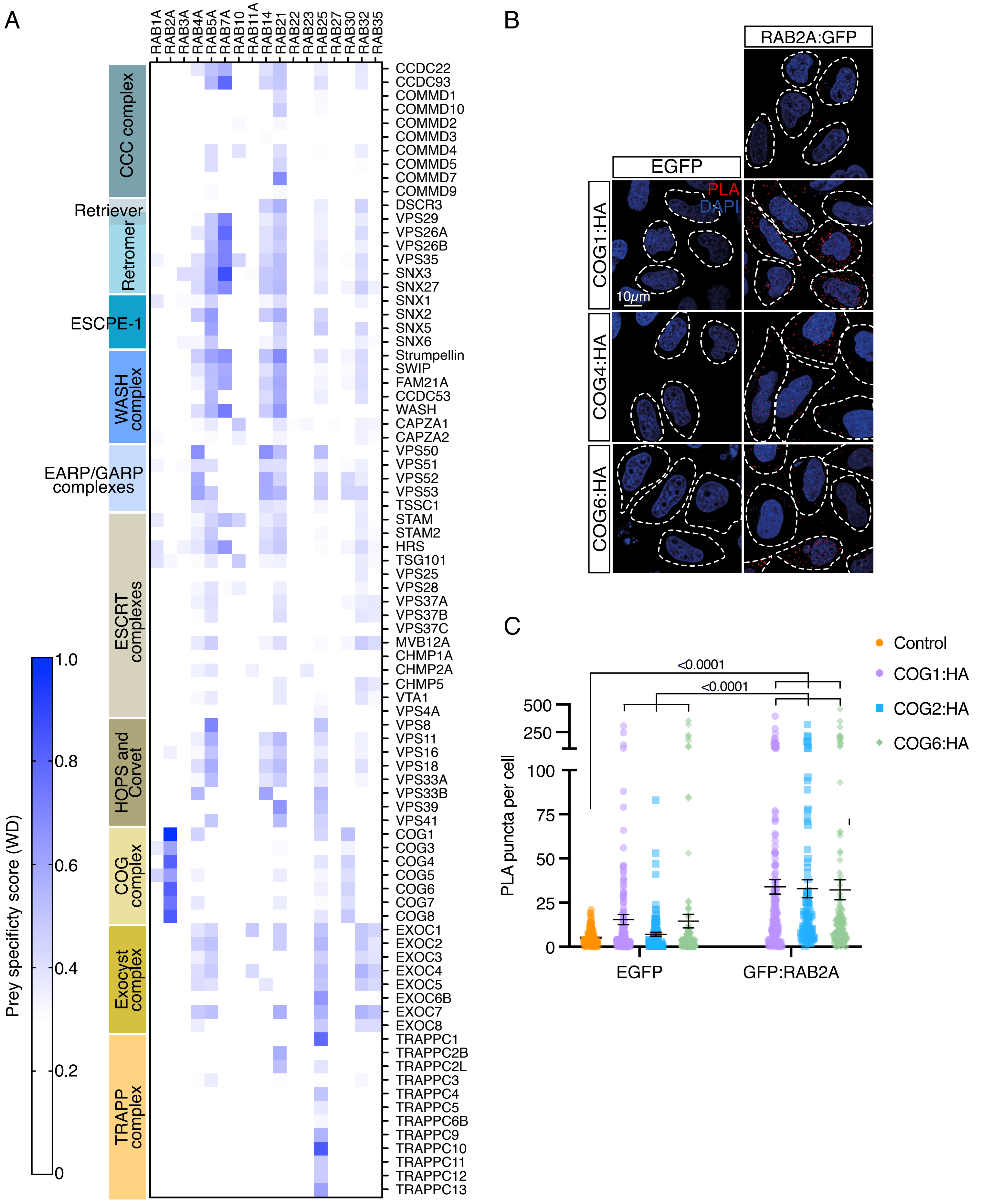

**Supplementary Figure 5**

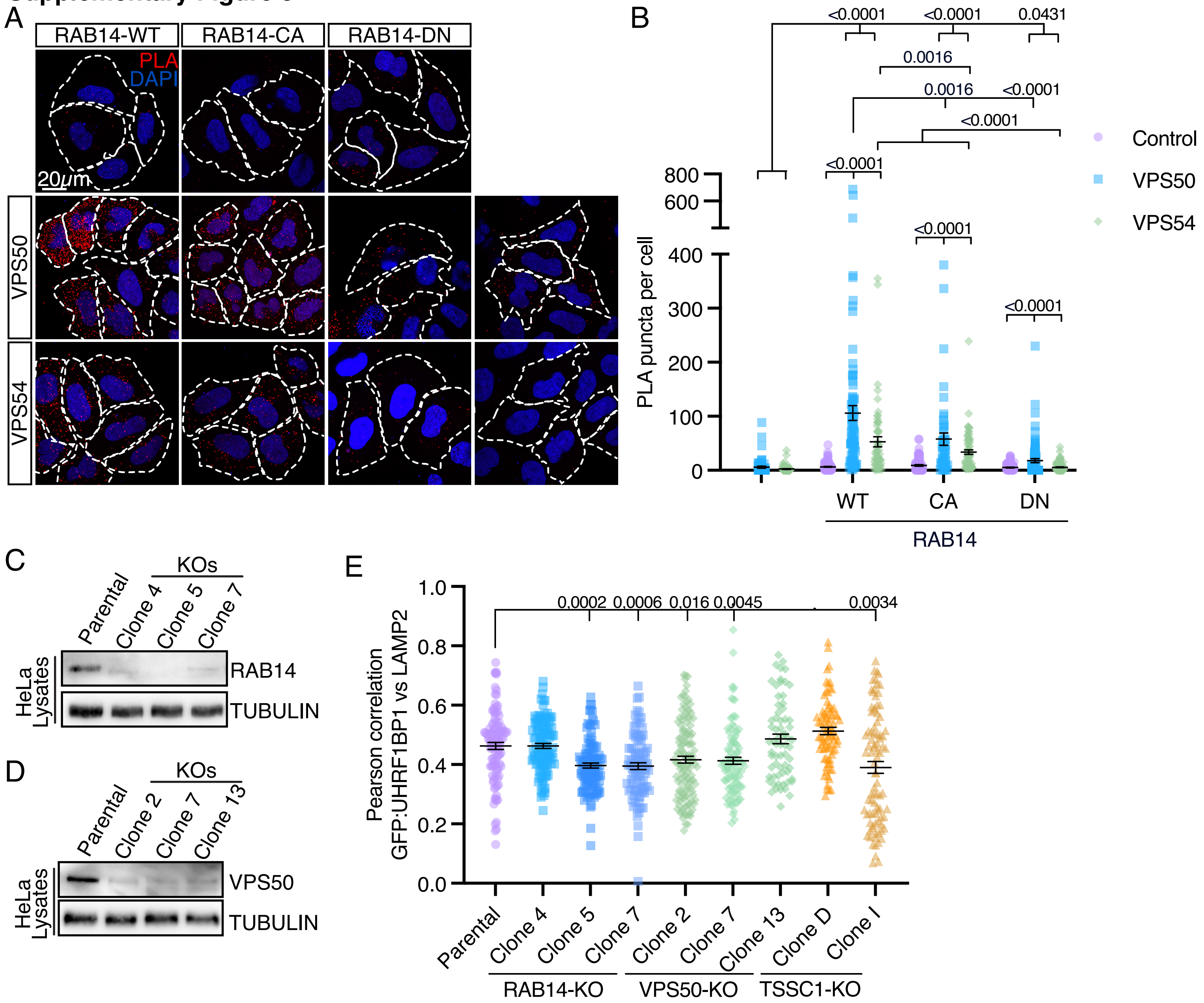
